# DeLUCS: Deep Learning for Unsupervised Clustering of DNA Sequences

**DOI:** 10.1101/2021.05.13.444008

**Authors:** Pablo Millán Arias, Fatemeh Alipour, Kathleen A. Hill, Lila Kari

**Affiliations:** School of Computer Science, University of Waterloo, Waterloo, ON, Canada; Department of Biology, University of Western Ontario, London, ON, Canada

## Abstract

We present a novel **De**ep **L**earning method for the **U**nsupervised **C**lustering of DNA **S**equences (DeLUCS) that does not require sequence alignment, sequence homology, or (taxonomic) identifiers. DeLUCS uses Frequency Chaos Game Representations *(FCGR)* of primary DNA sequences, and generates “mimic” sequence FCGRs to self-learn data patterns (genomic signatures) through the optimization of multiple neural networks. A majority voting scheme is then used to determine the final cluster assignment for each sequence. The clusters learned by DeLUCS match true taxonomic groups for large and diverse datasets, with accuracies ranging from 77% to 100%: 2,500 complete vertebrate mitochondrial genomes, at taxonomic levels from sub-phylum to genera; 3,200 randomly selected 400 kbp-long bacterial genome segments, into clusters corresponding to bacterial families; three viral genome and gene datasets, averaging 1,300 sequences each, into clusters corresponding to virus subtypes. DeLUCS significantly outperforms two classic clustering methods (*K*-means++ and Gaussian Mixture Models) for unlabelled data, by as much as 47%. DeLUCS is highly effective, it is able to cluster datasets of unlabelled primary DNA sequences totalling over 1 billion bp of data, and it bypasses common limitations to classification resulting from the lack of sequence homology, variation in sequence length, and the absence or instability of sequence annotations and taxonomic identifiers. Thus, DeLUCS offers fast and accurate DNA sequence clustering for previously intractable datasets.

## Introduction

Traditional DNA sequence classification algorithms rely on large amounts of labour intensive and human expert-mediated annotating of primary DNA sequences, informing origin and function. Moreover, some of these genome annotations are not always stable, given inaccuracies and temporary assignments due to limited information, knowledge, or characterization, in some cases. Also, since there is no taxonomic “ground truth,” taxonomic labels can be subject to dispute (see, e.g., [1–3]). In addition, as methods for determining phylogeny, evolutionary relationships, and taxonomy evolved from physical to molecular characteristics, this sometimes resulted in a series of changes in taxonomic assignments. An instance of this phenomenon is the microbial taxonomy, which recently underwent drastic changes through the Genome Taxonomy Database (GTDB) in an effort to ensure standardized and evolutionary consistent classification [4–6].

The applicability of existing classification algorithms is limited by their intrinsic reliance on DNA annotations, and on the “correctness” of existing sequence labels. For example, alignment-based methods crucially rely on DNA annotations indicating the gene name and genomic position. Similarly, supervised machine learning algorithms rely on the training data having stable taxonomic labels, since they carry forward any current misclassifications into erroneous future sequence classifications. To avoid these limitations, and given the ease of extensive sequence acquisition, there is a need for highly accurate unsupervised machine learning approaches to sequence classification that are not dependent on sequence annotations.

We propose a novel **De**ep **L**earning method for the **U**nsupervised **C**lustering of DNA **S**equences (DeLUCS), that is independent of sequence labels or annotations, and thus is not vulnerable to their inaccuracies, fluctuations, or absence. DeLUCS is, to the best of our knowledge, the first *highly-effective/light-preparation* DNA sequence clustering method, in that it achieves high “classification accuracies” while using only a minimum of data preparation and information. Indeed, the only information external to the primary DNA sequence that is used by DeLUCS is the implicit requirement that all sequences be of the same type (nuclear DNA, mtDNA, plastid, chloroplast), and that the selection of the dataset be based on some taxonomic criteria. Importantly, DeLUCS does not need any DNA annotations, does not require sequence homology or similarity in sequence lengths, and does not use any true taxonomic labels or sequence identifiers during the learning process.

In fact, DeLUCS only uses Frequency Chaos Game Representations with resolution *k* (*FCGR_k_*) of primary DNA sequences to find their cluster assignments, in an unsupervised manner. (In this paper, *k* = 6 was empirically selected as achieving the optimal balance between accuracy and time complexity.) This being said, the *post hoc* step of evaluating the performance of DeLUCS uses, by necessity, true taxonomic labels to quantify the accuracy of the computed clusters. This additional step is implemented via the Hungarian algorithm [7], which finds the optimal mapping between the numerical cluster labels determined by DeLUCS and the true taxonomic cluster labels. Based on this mapping, the resulting taxonomic cluster labels are then used to calculate the method’s accuracy (termed thereafter “classification accuracy”). Note that, for readability purposes, we will hereafter use true taxonomic labels for clusters, even though DeLUCS only outputs numerical cluster labels.

DeLUCS compensates for the absence of information external to the primary DNA sequence by leveraging the capability of deep learning to discover patterns (genomic signatures) in unlabelled raw primary DNA sequence data. DeLUCS is alignment-free, and learns clusters that match true taxonomic groups for large and diverse datasets, with high accuracy: 2,500 vertebrate complete mitochondrial genomes at multiple taxonomic levels, with accuracy ranging from 79% to 100%; 3,200 randomly selected bacterial genome segments, with an average length of 400 kbp, into bacterial families, with accuracy of 77% (inter-phylum) and 90% (intra- phylum); several datasets of viral gene sequences and of full viral genomes, averaging 1,300 sequences each, into virus subtypes, with accuracy of 99%, and 100% respectively. To the best of our knowledge, these are the largest real datasets classified to date, in clustering studies of genomic data: The biggest dataset analyzed in this paper totals over 1 billion bp of data, a full order of magnitude bigger than previous studies [8–14]. In addition, all but the viral gene dataset would be impossible to classify with alignment-based methods, due either to the prohibitive time cost of multiple sequence alignment or to the lack of sequence homology.

A direct comparison shows that DeLUCS significantly outperforms two classic algorithms for clustering unlabelled datasets (*K*-means++ and Gaussian Mixture Models, GMM), sometimes by as much as 47%. We also note that, for the majority of the computational tests, the DeLUCS classification accuracy is also comparable to, and sometimes higher than, that of a supervised machine learning algorithm with the same architecture.

DeLUCS is a fully-automated method that determines cluster assignments for its input sequences, independent of any homology or same-length assumptions, and oblivious to sequence taxonomic labels. DeLUCS can thus be used for successful *ab initio* classification of datasets that were previously unclassifiable by alignment-based methods, as well as datasets with uncertain or fluctuating taxonomy, where supervised machine learning methods are biased by their reliance on current taxonomic labels.

In summary, DeLUCS is the first effective alignment-free method that uses deep learning for unsupervised clustering of unlabelled raw DNA sequences. The main contributions of this paper are:

- DeLUCS clusters large and diverse datasets, such as complete mitochondrial genomes at several taxonomic levels; randomly selected bacterial genome segments into families; viral genes and viral full genomes into virus subtypes. To date, these are the largest real datasets of genomic data to be clustered, with our largest experiment comprising over 1 billion bp of data, a full order of magnitude larger than previous studies [8–14].
- DeLUCS achieves “classification accuracies” of 77% to 100%, in each case significantly higher than classic alignment-free clustering methods (K-means++ and GMM), with double digit improvements in most cases. For the majority of the computational tests, DeLUCS classification accuracies are also comparable to, or higher than, those of a supervised machine learning algorithm with the same architecture.
- DeLUCS is a highly-effective/light-preparation method for unsupervised clustering of DNA sequences. Its high classification accuracies are a result of combining the novel concept of mimic sequences with the invariant information learning framework and a majority voting scheme. It is termed light-preparation because it does not require sequence homology, sequence-length similarity, or any taxonomic labels/identifiers during the learning process.

### Prior Approaches

The time-complexity limitations of alignment-based methods, [15], in addition to their reliance on extraneous sequence information such as sequence homology, have motivated the development of numerous alignment-free methodologies [16, 17]. Of these, methods based on k-mer counts have been among the fastest and the most widely used [17]. In parallel to alignment-free approaches, machine learning methods have emerged as promising alternatives for solving classification problems both in genomics and biomedicine [18].

Fig 1 illustrates a summary of methods that combine alignment-free approaches with machine learning for genomic classification/clustering tasks. (The difference between classification and clustering is that, while in classification methods the cluster labels are given *a priori,* in clustering methods the clusters are “discovered” by the method.)

**Fig 1.**
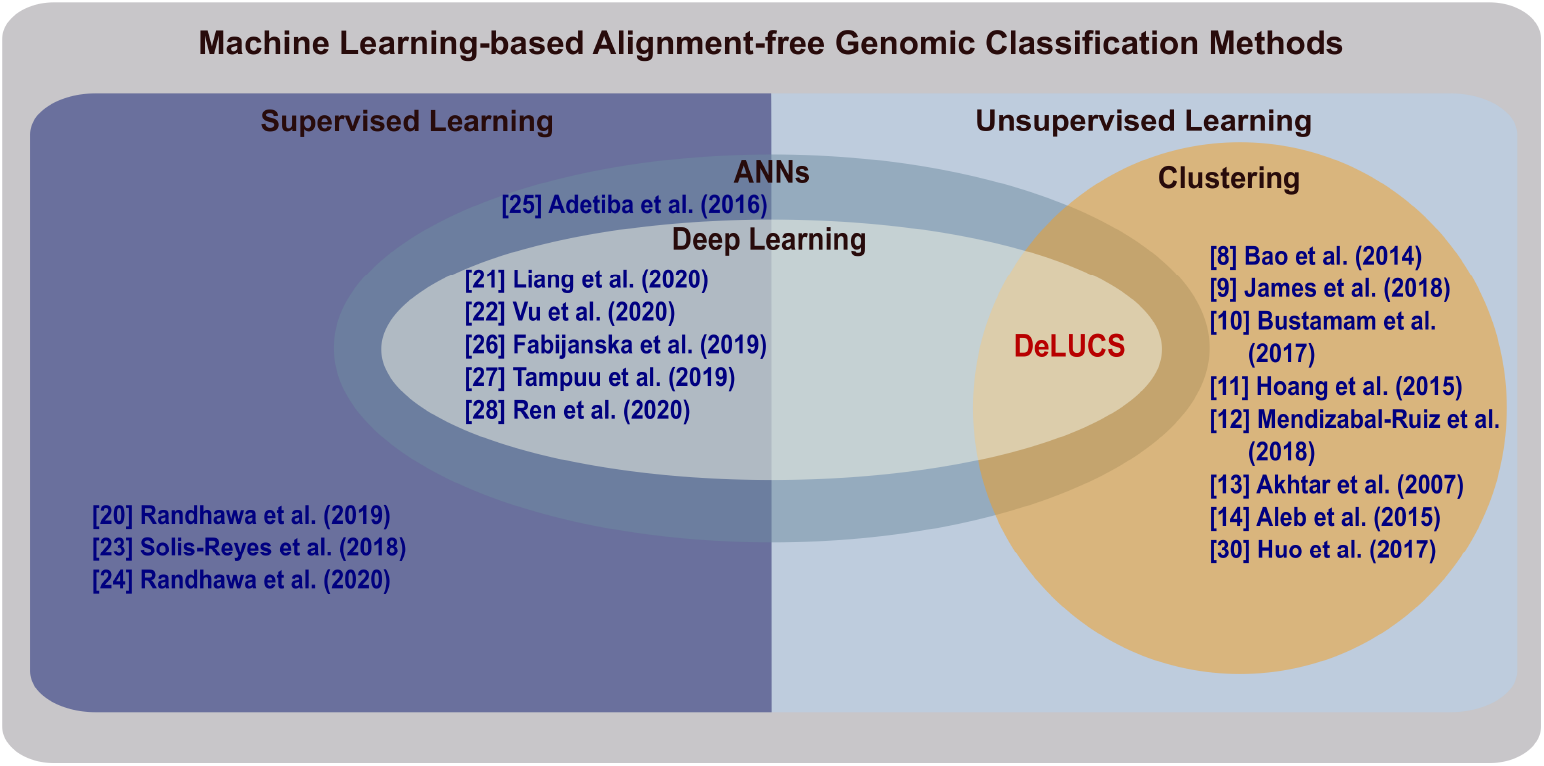
Machine learning-based alignment-free methods for classification/clustering of DNA sequences. DeLUCS is the first method to use deep learning for accurate unsupervised clustering of unlabelled raw DNA sequences. The novel use of deep learning in this context significantly boosts the classification accuracy (as defined in the Evaluation section), compared to two other unsupervised machine learning clustering methods (*K*-means++ and GMM).

#### Supervised Machine Learning Approaches

Among supervised learning algorithms, Artificial Neural Networks (ANNs) have proven to be the most effective, with ANN architectures with several layers of neurons (“deep learning”) being the top performers [19].

In the context of genome classification, alignment-free methods that employ supervised machine learning have been shown to outperform alignment-based methods in the construction of high-quality whole-genome phylogenies [20], profiling of microbial communities [21], and DNA barcoding at the species level [22]. In recent years, alignment-free methods have successfully applied supervised machine learning techniques to obtain accurate classification of HIV subtypes, [23], as well as accurate and early classification of the SARS-CoV-2 virus (COVID-19 virus) [24]. The increasing success of machine learning, and in particular deep learning, techniques is partly due to the introduction of suitable numerical representations for DNA sequences and the ability of the methods to find patterns in these representations (see [23, 25], respectively [21]). Other classification tasks in genomics such as taxonomic classification [26], and the identification of viral sequences among human samples from raw metagenomic segments [27, 28] have also been explored from the deep learning perspective.

One limitation of supervised deep learning is that the performance of the ANNs is heavily dependent on the number of labelled sequences that are available during training. This can become a limiting factor, even though raw sequencing data can now be obtained quickly and inexpensively, [29]. The reason for this is the intermediate process that lies between obtaining a raw DNA sequence and uploading that sequence onto a public sequence repository, namely the “invisible” work that goes into assigning a taxonomic label and attaching biological annotations. This is a laborious, expensive, and time consuming multistep process, comprising *ad hoc* wet lab experiments and protocols that cannot be automated due to the human expertize required. Another limitation of supervised learning is its sensitivity to perturbations in classification, since any present misclassifications in the training set are “learned” and propagated into future classification errors.

To overcome these limitations, one can attempt to use unsupervised learning, which operates with unlabelled sequences and compensates for the absence of labels by inferring identity-relevant patterns from unlabelled training data. Moreover, unsupervised learning does not perpetuate existing labelling errors, as the algorithms are oblivious to labels. It can correctly classify sequences of a type never seen during any previous training, by assigning the sequences to dynamically defined new clusters.

#### Unsupervised Machine Learning Approaches

Unlike supervised learning, in unsupervised learning training samples are unlabelled, i.e., the cluster label associated with each DNA sequence is not available (or is ignored) during training. In general, clustering large datasets using unsupervised learning is a challenging problem, and the progress in using unsupervised learning for clustering of genomic sequences has not been as rapid as that of its supervised classification counterparts. The effort made so far in the development of unsupervised alignment-free clustering algorithms for genomic sequences has been mainly focused on using generic clustering algorithms such as *K*-means or Gaussian Mixture Models (GMM) for different numerical representations of DNA sequences. For example, Bao et al., [8] used a representation of DNA sequences based on their word counts and Shannon entropy, whereby each sequence is represented by a 12-dimensional vector and the clustering is performed using *K*-means with Euclidean distance. James et al., [9] grouped DNA sequences based on four different similarity measures obtained from an alignment-free methodology that used *k*-mer frequencies and an adaptation of the mean shift algorithm, normally used in the field of image processing. Similar work [10,14,30] also builds on the *K*-means algorithm and *k*-mer counts. Another approach is the use of digital signal processing [11–13], whereby Fourier spectra calculated from a numeric representation of a DNA sequence are used as their quantitative description, and the Euclidean distance is used as a measure of dissimilarity to be employed by either the *K*-means or the GMM clustering algorithms.

Although *K*-means (and its improved version *K*-means++) is a simple and versatile algorithm, it is dependent on several restrictive assumptions about the dataset, such as the need for manual selection of the parameter *K*, and the assumption that all clusters have the same size and density. It is also heavily dependent on the selection of initial cluster centroids, meaning that for large datasets, numerous initializations of the centroids are required for convergence to the best solution and, moreover, that convergence is not guaranteed [31]. Although GMM is more flexible in regards to the distribution of the data and does not assume that all clusters are spherical, the initialization of clusters is still challenging, especially in high dimensional data, [32, 33].

A potential solution to these drawbacks could lie in recent developments in the field of computer vision [34–36], specifically in the concepts at the core of *invariant information clustering* (IIC) [36], one of the successful methods for the clustering of unlabelled images. These methods are effective for visual tasks and, as such, are not applicable to genomic data. In this paper, we propose the use of Frequency Chaos Game Representations (FCGR) of DNA sequences and the novel notion of *mimic sequences,* to leverage the idea behind IIC. In our approach, FCGR pairs of sequences and of their mimics are generated, and used as input for a *de novo* simple but general Artificial Neural Network (ANN) architecture, specifically designed for the purpose of DNA sequence clustering. Finally, majority voting over several independently trained ANN copies is used, to obtain the accurate cluster assignment of each sequence.

## Materials and Methods

In this section, we first give an overview of our method and the computational pipeline of DeLUCS. We then describe the core concepts of invariant information clustering, and detail how these concepts are adapted to DNA sequence clustering, by introducing the notion of “mimic sequences”. This is followed by a description of the architecture of the neural networks employed, the evaluation scheme used for assessing the performance of DeLUCS, and all of the implementation details. Finally we give a description of all the datasets used in this study.

### Method Overview

DeLUCS employs a graphical representation of DNA sequences introduced by Jeffrey in [37], called Chaos Game Representation (CGR). In this paper, we use a quantized version of CGR, called Frequency CGR with resolution *k*, and denoted by *FCGR_k_*. The *FCGR_k_* of a DNA sequence is a two-dimensional unit square image, with the intensity of each pixel representing the frequency of a particular k-mer in the sequence [38]. *FCGR_k_* is a compressed representation of the original DNA sequence, with the degree of compression indicated by the resolution *k*. All computational experiments in this paper use *k* = 6, which was empirically assessed as achieving the best balance between accuracy and time complexity. Several studies have demonstrated that the CGR of a genomic sequence can serve as its *genomic signature*, defined by Karlin and Burge [39] as any numerical quantity that is more similar for DNA sequences of closely related organisms, while being dissimilar for DNA sequences of more distantly related organisms, see Fig 2.

**Fig 2.**
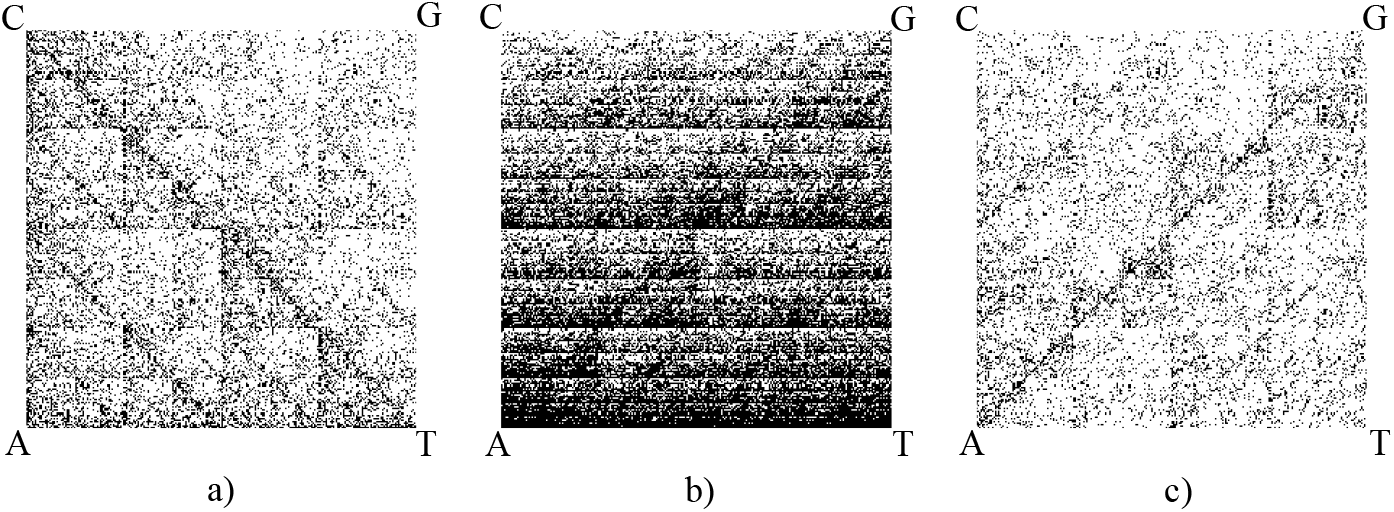
Chaos Game Representation of (a) the complete mitochondrial genome of *Rana Chosenica* (a frog), 18,357 bp – Accession ID: NC_016059.1; (b) the first 80,000 bp of the *Bacillus mycoides* genome – Accession ID: NZ_CP009691.1; (c) the complete genome of *Dengue Virus 2,* 10,627 bp – Accession ID: GU131948.1

Unlike some quantization techniques [40, 41], DeLUCS does not need an intermediate step of supervised learning to produce a compressed representation of data, retaining only the information needed for the correct classification of the feature vector and the cluster assignment. This is because *FCGR_k_* already is a compressed representation, by virtue of storing only the counts of all *k*-mers in the sequence for a given value of *k*. These *k*-mer counts contain the intrinsic, taxonomically relevant, information used for the unsupervised learning in DeLUCS.

The general pipeline of DeLUCS, illustrated in Fig 3, consists of three main steps:

1. For each DNA sequence in the dataset several artificial *mimic sequences* are constructed, and considered to belong to the same cluster. These mimic sequences are generated using a probabilistic model based on transversions and transitions. The *k-mer* counts for both the original sequence and its mimic sequences are then computed, to produce their respective *FCGRs.* In this study, *k* = 6 was empirically assessed as achieving the best balance between high accuracy and speed.
2. Pairs consisting of the FCGR of the original DNA sequence and the FCGR of one of its mimic sequences are then used to train several copies of an Artificial Neural Network (ANN) independently, by maximizing the mutual information between the network predictions for the members of each pair.
3. As the training process of the ANNs is a randomized algorithm which produces different outcomes with high variance, a majority voting scheme over the outcomes of the ANNs in Step 2 is used to determine the final cluster assignment for each sequence.

**Fig 3.**
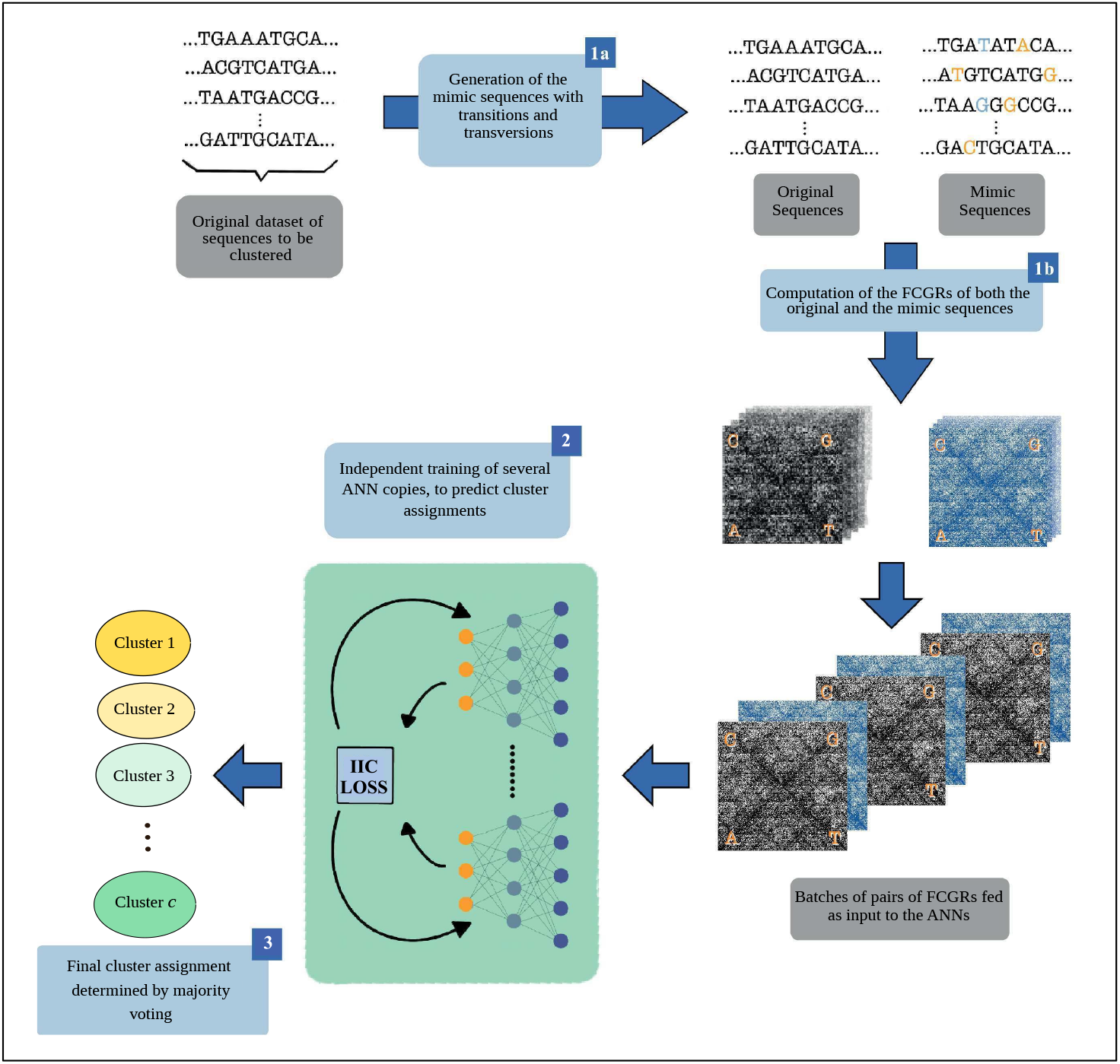
General DeLUCS pipeline. The input consists of the original DNA sequences to be clustered. (1a): Artificial *mimic sequences* are generated from the original sequences, by using a probabilistic model based on transitions and transversions. (1b): FCGRs of all original (black) sequences, and of all mimic (blue) sequences are computed, and data pairs of the form *“FCGR of DNA sequence, FCGR of one of its mimics”* are divided in batches for the training process. (2): Several copies of the ANN are trained independently, with the loss function being the negative mutual information between the network predictions for a sequence and that of its mimic. (3): Majority voting is used to obtain the final cluster assignment for each sequence.

To evaluate the quality of the clusters, an additional step is performed, independent of DeLUCS. This step first utilizes the Hungarian algorithm to determine the optimal correspondence between the cluster assignments learned by DeLUCS and the true taxonomic cluster labels. It then proceeds to determine the accuracy of the DeLUCS cluster predictions, as detailed in the Evaluation section.

### Invariant Information Clustering (IIC)

Steps 1 and 2 in the DeLUCS pipeline build upon the underlying concepts of IIC, [36], which leverages some information theory notions described in this subsection.

Given a discrete random variable *X* that takes values *x* ∈ *χ* and has probability mass function *p*(*x*) = *P*(*X = x*), the entropy *H*(*X*) is a measure of the average uncertainty in the random variable and is defined by

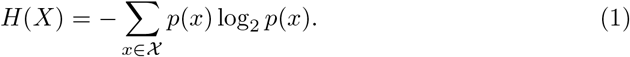

*H*(*X*) also represents the average number of bits required to describe the random variable *X*.

Given a second random variable 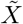 that takes values 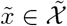, we can also define the conditional entropy 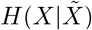, for a pair sampled from a joint probability distribution 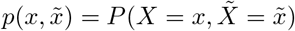, as the entropy of a random variable *X* conditional on having some knowledge about the variable 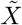. The reduction in the uncertainty of *X* introduced by the additional knowledge provided by 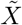 is called *mutual information* and it is defined by

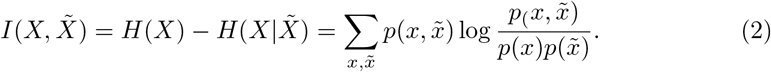

The mutual information measures the dependence between the two random variables, and it represents the amount of information that one random variable contains about another. 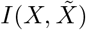 is symmetric, always non-negative, and is equal to zero if and only if *X* and 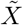 are independent.

The information bottleneck principle [42, 43], which is part of the information theoretic approach to clustering, suggests that clusters only need to capture relevant information. In order to filter out irrelevant information, IIC aims to learn only from paired data, i.e., from pairs of samples 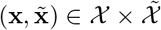 taken from a joint probability distribution 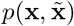. If, for each pair, 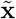 is an artificially created copy of **x**, it is possible to find a mapping Φ that encodes what is common between **x** and 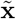, while dropping all the irrelevant information. If such a mapping Φ is found, the image 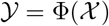 becomes a compressed representation of the original space *χ*.

To find the best candidate for Φ, one way is to make Φ(**x**) represent a random variable, and then maximize the predictability of sample **x** from sample 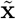 and vice versa, that is, find a mapping Φ(**x**) that maximizes 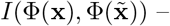 the mutual information between the encoded variables – over all **x** ∈ **χ**.

This idea suggests that Φ can be calculated using a deep neural network with a *softmax* as the output layer. For a dataset with an expected number of *c* clusters, *c* ∈ ℕ, the output space will be 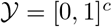, where for each sample **x** we have that Φ(**x**) represents the distribution of a discrete random variable over the *c* clusters. The mutual information can be modified with the introduction of a hyper-parameter λ ∈ ℝ that weighs the contribution of the entropy term in Eq (2). However, instead of maximizing the weighted mutual information, we use a numerical optimizer to minimize its opposite (mathematically, the negative weighted mutual information) during the training process of the ANN. Hence, the loss function to be minimized becomes:

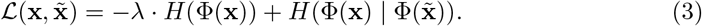

In Eq (3), the entropy term *H*(Φ(**x**)) measures the amount of randomness present at the output of the network, and it is desirable for that value to be as large as possible, in order to prevent the architecture from assigning all samples to the same cluster. The conditional entropy term 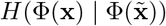 measures the amount of randomness present in the original sample **x**, given its correspondent 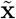. This conditional entropy should be as small as possible, since the original sample **x** should be perfectly predictable from 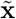.

### Generation of Mimic Sequences

The success of the method described in the previous section is fundamentally dependent on the way 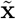 is artificially generated from **x**. In the particular case of our application, where the samples **x** are DNA sequences, we refer to the artificially created 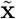 as *mimic sequences* (sometimes called simply mimics). In this context, the generation of mimic sequences poses the additional challenge that they should be sufficiently similar to the originals so as not to be assigned to a different cluster.

Given a set *X* = {*x*_1_,..., *x_n_*} of *n* DNA sequences, we construct the set of pairs

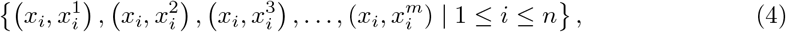

where *m* ≥ 3 is a parameter representing the number of mimic sequences generated for each original sequence *x_i_*, 1 ≤ *i* ≤ *n*. We use a simple probabilistic model based on DNA substitution mutations (transitions and transversions) to produce different mimic sequences, as follows. Given a sequence *x_i_* and a particular position *j* in the sequence, we fixed independent transition and transversion probabilities *p_ts_* [*j*] and *p_tv_* [*j*] respectively. Next, we produce the following mimic sequences, probabilistically: 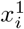 with only transitions, 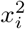 with only transversions, and 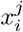 with both transitions and transversions, for all 3 ≤ *j* ≤ *m*. The parameter *m* is determined, for each experiment, based on the particulars of its dataset. Its default value is 3, to account for the use of the two individual substitution mutations and their combination, but may have to be increased if the number of available sequences per cluster is insufficient to obtain a high classification accuracy.

The rationale behind using transition and transversion probabilities to generate sequence mimics is biologically inspired. That being said, we use this method only as a mathematical tool without attributing any biological significance, to create minimally different sequences through randomly distributed base substitutions. In this paper we use probabilities *p_ts_* = 10^-4^ and *p_tv_* = 0.5 × 10^-4^, assessed empirically to result in the best classification accuracies. Although the mutation rates used are biologically inspired, they are not biologically precise given that mutation rates vary regionally, with species [44, 45], and with the estimation method [46]. Lastly, in practice, with no taxonomic label, it is impossible to select species-specific mutation rates.

### Artificial Neural Network (ANN) Architecture

The pairs of FCGRs of the original DNA sequences and their mimic sequences are used as inputs, to train several independent copies of an ANN. Since the size of the genomic datasets under study is at least an order of magnitude smaller than what is used in computer vision, we noted that the common architectures that have proven effective in the application of deep learning for various visual tasks were not suitable for our datasets. Hence, we designed, *de novo*, a simple but general architecture that is suitable for the clustering of DNA sequences. The complete architecture is presented in Fig 4 and it consists of two fully connected layers, *Linear (512 neurons)* and *Linear (64 neurons)*, each one followed by a Rectified Linear Unit (*ReLU*) and a *Dropout* layer with dropout rate of 0.5. The output layer *Linear* (*c_clusters*), where *c* is a numerical parameter representing the upper bound of the number of clusters, is followed by a *Softmax* activation function.

**Fig 4.**
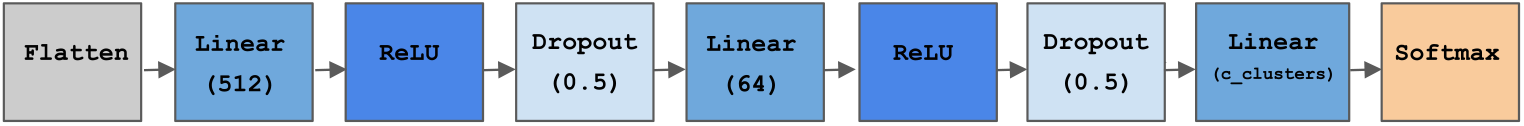
Architecture of the deep Artificial Neural Network used in this paper. The input FCGRs are flattened into one-dimensional representations prior to entering the first *Linear* layer. The parameter in each linear layer except the output layer represents the number of neurons. For the *Dropout* layer, the parameter represents the dropout rate. The parameter c (in *c_clusters)* of the output linear layer, represents the expected upper bound of the number of clusters. The *Softmax* layer is used to obtain a probability distribution as the output of the network.

The network receives as input pairs of FCGRs (two-dimensional representations of DNA sequence composition) and flattens them into one-dimensional representations, which are then fed sequentially to the first Linear layer. The inclusion of ReLUs is essential for the training process, as they help mitigate the problems of vanishing gradients and other back-propagation errors. The dropout layers prevent the model from over-fitting, which in unsupervised learning comes in the form of degenerate solutions, i.e., all the samples being assigned to the same cluster. Finally, the Softmax layer gives as output a *c*-dimensional vector Φ(**x**) ∈ [0, 1]^*c*^, such that Φ_c_j__ (**x**), 1 ≤ *c_j_* ≤ *c*, represents the probability that an input sequence **x** belongs to a particular cluster *c_j_*.

Note that this general architecture was designed so as to be successful for the clustering of all the diverse datasets presented in this study. However, the main pipeline of DeLUCS allows it to be used also with other architectures, including architectures that make use of the two-dimensional nature of the FCGR patterns and are performant for specific types of genomic data (e.g., Convolutional Neural Networks).

### Evaluation

Clustering results can be evaluated using both internal and external validation measures. Internal validation methods [47, 48] evaluate clustering algorithms by measuring some of the discovered clusters’ internal properties (e.g., separation, compactness), while external validation methods [49–51] evaluate clustering algorithms by measuring the similarity of the discovered clustering with the ground truth. We note that many genomic datasets are sparse, incomplete, and subject to sampling bias (more than 86% of existing species on Earth and 91% of species in the oceans have not yet been classified and catalogued [52]). Thus, when taxonomic ground truth is available for an external validation, agreement between discovered clusters and real taxonomic groups is preferable to, and more informative than, internal validation methods.

We include performance comparison results obtained using (unsupervised) classification accuracy (ACC), as ACC uses the optimal mapping between the discovered clusters and the ground truth clusters, and has been used extensively in recent deep unsupervised learning studies [53]. Comparison results obtained using two other external evaluation methods, normalized mutual information of the partitions (NMI), and adjusted rand index (ARI), lead to similar conclusions, and can be found in *S6 Appendix: Using NMI and ARI to compare DeLUCS with K-means++ and GMM.*

In calculating the classification accuracy ACC, we follow the standard protocol that uses the confusion matrix as the cost matrix for the Hungarian algorithm [35, 36], to find the optimal mapping *f* that assigns to each cluster label *c_j_*, 1 ≤ *c_j_* ≤ *c* found by DeLUCS, a taxonomic label *f* (*c_j_*). We then use this optimal assignment to calculate the classification accuracy of the unsupervised clustering method, which is defined as:

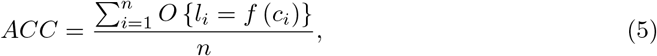

where *n* is the total number of sequences, and for each DNA sequence *x_i_*, 1 ≤ *i* ≤ *n*, we have that is its true taxonomic label, *c_i_* is the cluster label found by the algorithm, *f* (*c_i_*) is the taxonomic label assigned to *c_i_* by the optimal mapping *f*, and *O* is a comparison operator returning 1 if the equality in the argument holds and 0 otherwise.

Several unsupervised clustering methods exist in the literature, and various deep learning-based clustering tools have been adapted to bioinformatics [53]. However, these methods are domain specific and are not optimized to operate with DNA sequences, hence they perform poorly when used with DNA datasets (see *S5 Appendix: A note on comparing DeLUCS with other deep learning-based clustering methodologies).* In this paper, K-means++ and Gaussian Mixture Model (GMM) were selected for comparison with DeLUCS because they are general clustering algorithms at the core of many unsupervised learning frameworks, and have been previously used with DNA sequence datasets [8,10,13,14]. Note that the Hungarian algorithm is used to find the mapping that maximizes ACC for all the unsupervised clustering methods considered. In all three cases, the use of the true taxonomic labels is for *post hoc* evaluation purposes only, as true labels are never used during the training process.

Lastly, we compare these three unsupervised clustering methods to a supervised learning classification method. For this purpose, the same neural network architecture described in the previous section is trained, using labelled data and the cross-entropy loss function. The accuracy of the classification is calculated by first taking 70% of the data for training, and 30% of the data for testing. The classification accuracy of the supervised learning method is then defined as the ratio of the number of correctly predicted testing sequence labels to the total number of testing sequences.

### Implementation

During the training procedure, all the hyperparameters of the method are fixed and common to all of the tests, and were empirically selected as yielding the best performance. All the flattened *FCGR_k_* were normalized before being fed into the network, by using the *L*1 norm, i.e., by dividing the values of each *k*-mer count vector by their sum. This normalization brings all of the inputs of the ANN into the same range of values, which contributes to the reliability of the ANN convergence.

The networks are initialized using the Kaiming method [54], to avoid exponential reduction or magnification of the input magnitudes. This is crucial for our method because a poor initialization may lead to degenerate solutions, as one of the terms in the loss function becomes dominant. We use the Adam optimizer, [55], with a learning rate of 5 × 10^-5^, and the networks were trained for 150 epochs with no early stopping conditions. Another vital consideration during training is the selection of the batch size (empirically determined to be 512), because the marginalization that is performed to find the distribution of the output is done over each batch of pairs. If the batch size is not large enough to represent the real distribution of the data, the entropy term in the loss function becomes dominant, leading to sub-optimal solutions. Lastly, we fix the value of the hyperparameter λ to 2.5 (in Eq 3).

DeLUCS is fully implemented in Python 3.7, and the source code is publicly available in the Github repository https://github.com/pmillana/DeLUCS. Users may reproduce the results obtained in this paper, or use their own datasets for the purpose of clustering new sequences (see *S1 Appendix: Instructions for reproduction of the tests using DeLUCS).* All of the tests were performed on one of the nodes of the Cedar cluster of Compute Canada (2 x Intel E5-2650 v4 Broadwell @ 2.2GHz CPU, 32 GB RAM) with NVIDIA P100 Pascal(12G HBM2 memory).

### Datasets

We used three different datasets in this study to confirm the applicability of our method to different types of genomic sequences (mitochondrial genomes, randomly selected bacterial genome segments, viral genes, and viral genomes), and all data was retrieved from publicly available databases. Tables 1, 2 and 3, summarize the dataset details for each of the 11 computational tests performed (see *S2 Appendix: Query options for data download,* for more information).

**Table 1.** Details of the datasets used in computational tests 1 through 6 (full vertebrate mitochondrial genomes).

| Test # | Dataset | Total no.of seq. | Min clus. size | Max clus. size | Min. seq.len. (bp) | Avg. seq.len. (bp) | Max seq.len. (bp) |
| --- | --- | --- | --- | --- | --- | --- | --- |
| 1 | Subphylum Vertebrata (Actinopterygii (Fish): 500, Amphibians: 500, Birds: 500, Mammals: 500, Reptiles: 500) | 2,500 | 500 | 500 | 14,127 | 16,951 | 24,317 |
| 2 | Class Actinopterygii (Neopterygii: 40, Polypteriformes: 33, Chondrostei: 40) | 113 | 33 | 40 | 15,531 | 16,623 | 18,062 |
| 3 | Subclass Neopterygii (Ostariophysi: 250, Clupeomorpha: 250, Elopomorpha: 226, Acanthopterygii: 250, Paracanthopterygii: 249, Protacanthopterygii: 250) | 1,475 | 226 | 250 | 15,564 | 16,688 | 19,801 |
| 4 | Superorder Ostariophysi (Cypriniformes: 130, Characiformes: 123, Siluriformes: 130) | 383 | 123 | 130 | 15,664 | 16,635 | 17,998 |
| 5 | Order Cypriniformes (Cyprinidae: 80, Cobitidae: 80, Balitoridae: 75, Nemacheilidae: 80, Xenocyprididae: 80, Acheilognathidae: 70, Gobionidae: 80) | 545 | 70 | 80 | 16,061 | 16,610 | 17,282 |
| 6 | Family Cyprinidae (Acheilognathus: 47, Acrossocheilus: 46, Carassius: 45, Labeo: 45, Microphysogobio: 35, Notropis: 26, Onychostoma: 29, Rhodeus: 28, Schizothorax: 31, Sarcocheilichthys: 45, Cyprinus: 43, Sinocyclocheilus: 27) | 447 | 26 | 47 | 16,070 | 16,632 | 17,426 |

The choice of computational Tests 1 to 6 (datasets within the sub-phylum Vertebrata) follows a decision-tree approach whereby, at each taxonomic level, one particular cluster is selected for further in-depth analysis. Note that, since the minimum and maximum cluster sizes are different for each test, the number of sequences selected from a given taxon can differ from one test to another. For example, in Test 3 only 250 out of the total 2,723 available Ostariophysi sequences were selected, due to the need to achieve cluster balance, while the min/max cluster size parameters of Test 4 allowed for 383 Ostariophysi sequences to be used. The remaining Ostariophysi sequences could not be selected in either test since most of them belonged to the over-represented Order Cypriniformes (2,171 available sequences). Test 1 and 2 illustrate a different scenario: 500 of the total 7,876 available Actinopterygii sequences were used in Test 1, since this cluster size was sufficient to achieve high accuracy, while in Test 2 only 113 Actinopterygii sequences could be used, due to the under-representation of class Polypteriformes (33 available sequences) and over-representation of class Neopterygii (7,715 available sequences).

**Table 2.**
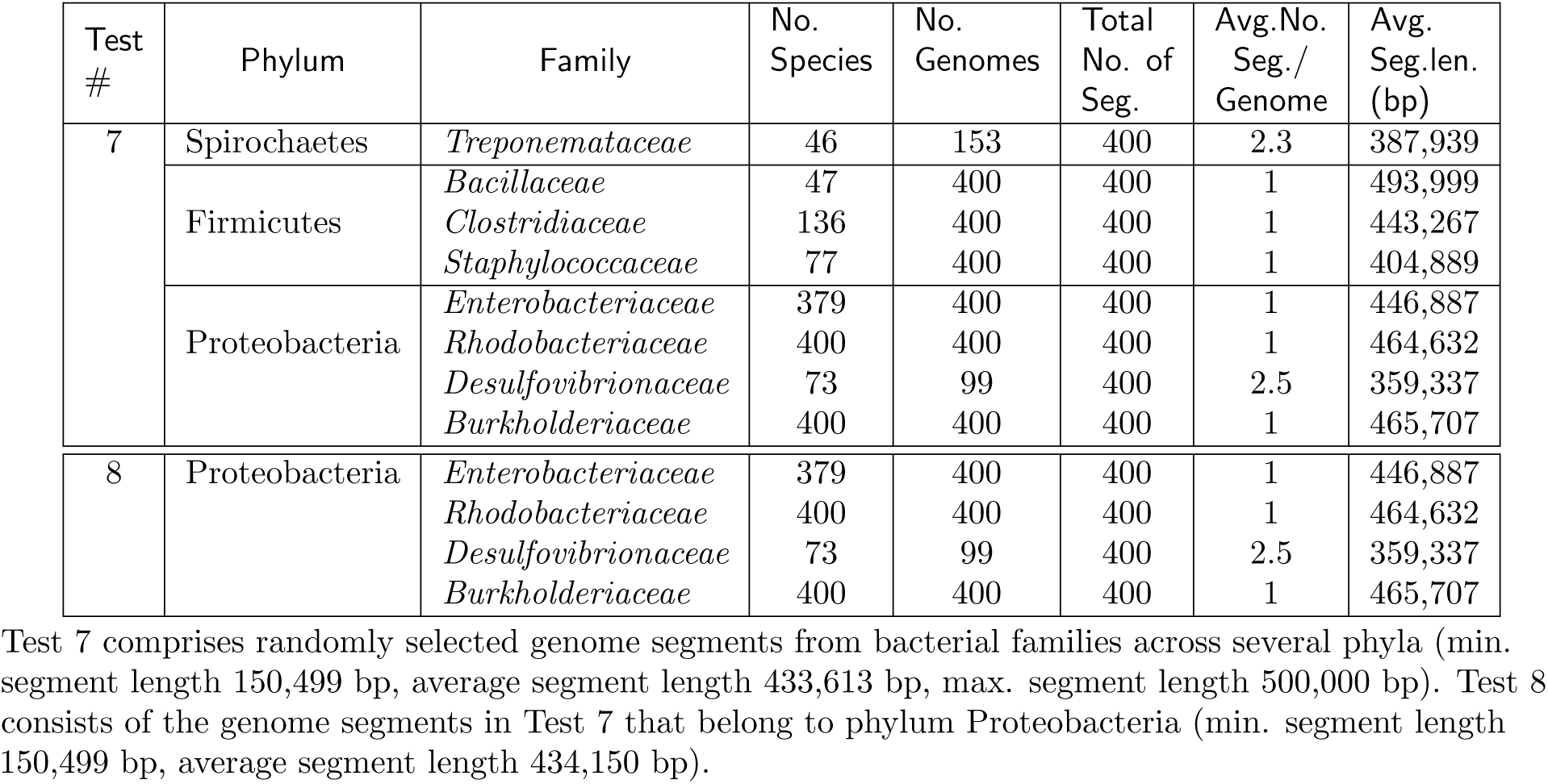
Details of the datasets used in computational test 7 and 8 (randomly selected bacterial genome segments, 400 segments per family, each of length between 150 kbp and 500 kbp).

**Table 3.** Details of the datasets used in computational tests 9, 10 and 11 (*Influenza virus* NA-encoding gene, *Dengue virus* full genomes, *Hepatitis B virus* full genomes).

| Test # | Dataset | Total no.of seq. | Min clus. size | Max clus. size | Min. seq.len. (bp) | Avg. seq.len. (bp) | Max seq.len. (bp) |
| --- | --- | --- | --- | --- | --- | --- | --- |
| 9 | Influenza A (NA-encoding gene)<br>(Subtypes H1N1: 191, H2N2: 187, H5N1: 188, H7N3: 193, H7N9: 190) | 949 | 187 | 193 | 1,345 | 1,409 | 1,469 |
| 10 | Dengue complete genomes<br>(Subtypes 1: 409 , 2: 409, 3: 408, 4: 407) | 1,633 | 407 | 409 | 10,161 | 10,559 | 10,991 |
| 11 | Hepatitis B complete genomes<br>(Subtypes A: 258, B: 262, C: 263, D: 260, E: 261, F: 258) | 1,562 | 258 | 263 | 3,182 | 3,210 | 3,227 |

#### Mitochondrial genomes (Table 1)

The first dataset consists of complete vertebrate mitochondrial genomes. We used the software Geneious 2020.2.4 to obtain a list with 13,300 accession numbers. Sequences were downloaded from the National Center for Biotechnology Information (NCBI) on November 16, 2020, after removing all of the sequences shorter than 14,000 bp and longer than 24,500 bp. For this dataset, the goal was to assess the performance of DeLUCS at different taxonomic levels, starting at the sub-phylum level, down to the genus level. To do so, the cluster with the largest number of sequences available at each taxonomic level was selected for the next computational experiment, to ensure that we could reach as deep as possible into the phylogenetic tree. A decision tree illustrating the cluster choices at different taxonomic levels is included in Supplementary Material (see *S4 Appendix: Detailed description of the mtDNA datasets,* and Fig 1 therein).

DeLUCS reaches optimal performance (in terms of classification accuracy) when a dataset consists of balanced clusters, that is, clusters that are similar in size, and when each cluster has more than a minimum number of sequences (herein 20 sequences). These two requirements, together with the number of sequences per cluster available on NCBI, were used to determine the minimum and maximum cluster size for each test.

After determining the minimum cluster size for a test (the size of the smallest cluster larger than 20), the clusters that were smaller than this test minimum were discarded. In addition, sequences that belonged to the parent taxon, but lacked a sub-taxon identifier (cluster label) were excluded (see *S4 Appendix: Detailed description of the mtDNA datasets,* for details).

Some clusters were larger than the test maximum (a number bigger than – but not more than 20% bigger than – the test minimum, to ensure cluster balance). For such clusters, sequences were randomly selected until the test maximum was reached. Note that since the minimum and maximum cluster size are test dependent, the number of sequences in a cluster is also test dependent (for example, Test 1 uses 500 Actinopterygii sequences, while Test 2 uses only 113). Moreover, for similar reasons, the number of sequences per cluster is usually smaller than the total number of sequences with that cluster label that are available on NCBI.

#### Bacterial genomic sequences (Table 2)

The second dataset consists of 3,200 randomly selected genomic segments of bacterial DNA from the eight different families analyzed in [11]. These bacterial families belong to three different phyla: *`Spirochaetes` (Treponemataceae*), *Firmicutes (Bacillaceae, Clostridiaceae,* and *Staphylococcaceae*), and *`Proteobacteria` (Enterobacteriaceae, Rhodobacteriaceae, Desulfovibrionaceae,* and *Burkholderiaceae*).

To construct a balanced dataset that captured as much diversity as possible, we considered all of the available species per family, according to GTDB (release 95), [56], and first excluded those species for which none of the sequences had a contig that was of the minimum length (herein, 150 kbp). For these computational tests, the cluster size was selected to be 400 sequences per cluster. This led to the following cases and corresponding experiment design choices:

1. The number of available species in a family is larger than 400 (*Rhodocacteriaceae* and *Burkholderiaceae*) We selected 400 species at random and, for each of the selected species, we randomly selected one genome. Then we selected a random segment of at most 500 kbp from each genome. If there was only one genome available for a particular species, and the length of its largest contig was between 150 kbp and 500 kbp, the entire contig was selected. The selection strategy was designed to include as many families as possible in the final dataset.
2. The number of available species in a family is less than 400

A. The total number of genomes in that family is larger than 400 (*Bacillaceae*, *Clostridiaceae*, *Staphylococcaceae*, *Enterobacteriaceae*) We calculated the median *M* of the number of genomes per species, for that family, and randomly chose at most *M* genomes per species (*M* = 2 for *Bacillaceae*, *M* = 1 for *Clostridiaceae*, *M* = 7 for *Staphylococcaceae*, and *M* = 2 *Enterobacteriaceae).* We then selected a random segment of at most 500 kbp from each genome. If there were fewer than M genomes representing a species, and the length of their largest contig was between 150 kbp and 500 kbp, then the entire contig was selected. Since after this selection more sequences were still required to reach the required 400 sequences per cluster, we chose the rest of the genomes at random from the family, without repetition, and selected a segment of 500 kbp from each such genome.
B. The total number of genomes in the family is less than 400 (*Treponemataceae* and *Desulfovibrionaceae)* For every genome, whenever possible, contigs were divided into successive segments of at most 500 kbp, that were added to a “pool”. If the length of a contig was between 150 kbp and 500 kp, the entire contig was added to the pool. From this pool, 400 segments were selected for inclusion in the final dataset. This selection strategy ensured that all of the species were represented, and that the shortest segments were selected last.

A description of the final composition of each cluster is presented in Table 2. In addition to the inter-phylum classification of bacterial sequences into families (Test 7), we assessed the performance of DeLUCS for an intra-phylum classification into families, within the `Proteobacteria` phylum only (Test 8). The dataset for Test 8 was simply the subset of the dataset in Test 7 that included only the segments from genomes in bacterial families from phylum `Proteobacteria`.

#### Viral genomes (Table 3)

The third group of datasets consists of three different sets of viral genome sequences. The minimum and maximum cluster sizes were determined in a manner similar to that of the mitochondrial DNA datasets, i.e., for all the tests, the minimum cluster size was fixed to the size of the smallest cluster available and the maximum cluster size was fixed manually to a number exceeding the minimum cluster size by no more than 20%, to ensure cluster balance.

To asses the performance of DeLUCS at the gene level, the dataset of Test 9 comprises 949 sequences of segment 6 of the *Influenza A virus* genome (encoding the neuraminidase protein, average length 1,409 bp). The sequences were downloaded from NCBI, [57], as belonging to subtypes H1N1, H2N2, H5N1, H7N3, and H7N9, as per [11]. The dataset of Test 10 comprises 1,633 *Dengue virus* full genome sequences downloaded from NCBI, [58], comprising four different subtypes. Finally, the dataset for Test 11 comprises 1,562 *Hepatitis B virus (HBV)* full genome sequences, downloaded from the Hepatitis Virus Database, [59], comprising six different subtypes. A description of the composition of each dataset is presented in Table 3.

## Results

### Qualitative and Quantitative DeLUCS Performance Results

We first used a qualitative measure to assess DeLUCS’s ability to group DNA sequences into meaningful clusters within the previously described datasets. Fig 5 illustrates how, during the training stage, the ANN discovers meaningful clusters as the learning process evolves. The progress of DeLUCS is demonstrated in terms of the number of epochs (one epoch means a single pass of the ANN through the entire dataset, trying to discover new patterns). In Fig 5, the number of clusters is *c* = 5 and it corresponds to the number of vertices in the regular polygon, whereby each vertex represents a taxonomic label. The coordinates of each point are calculated using a convex combination of the components of the c-dimensional probability vector (see section Artificial Neural Network (ANN) Architecture), [36]. When the learning process starts, all the sequences are located at the center of the polygon, which means that each sequence is equally likely to be assigned any of the five cluster labels. As the learning process continues, the network starts assigning the sequences to clusters (with similar sequences being grouped together closer to their respective vertex/cluster), with higher and higher probability. Note that if two sequences are assigned the same probability vectors, their corresponding points in Fig 5 will overlap.

**Fig 5.**
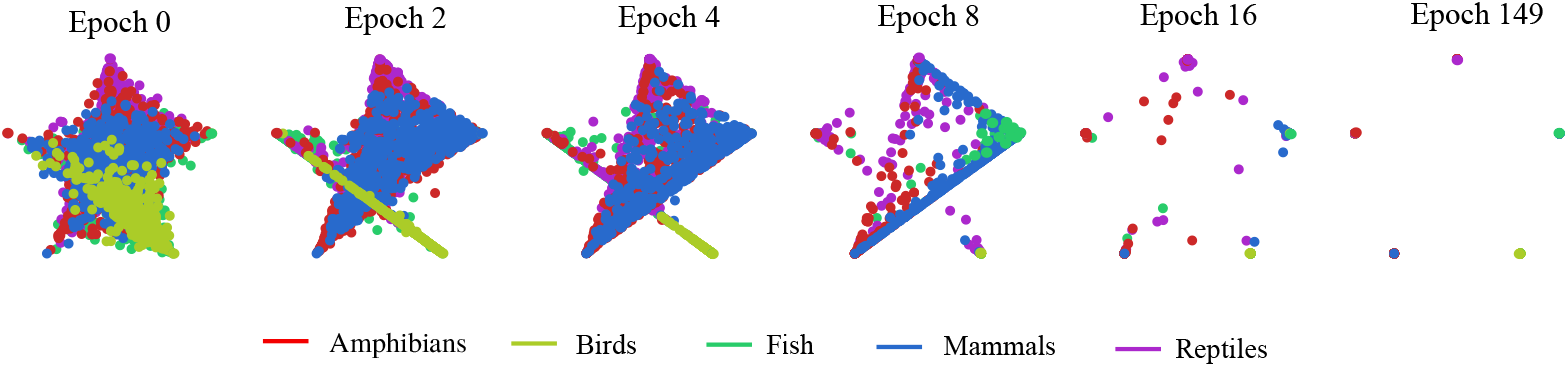
Learning process for the clustering of 2,500 vertebrate mtDNA full genomes into *c* =5 different clusters, each corresponding to one corner of the pentagon. Each point represents a sequence, and its position indicates the probability that it is assigned to different classes. Hence, a point in the center has equal probabilities to be assigned to all 5 vertices/clusters, while a point located in a vertex has probability of 1 of belonging to that vertex/cluster. With successive epochs, the learning progresses until, at epoch 149, all sequences are correctly placed in their respective vertex/cluster. Note that points in the figure overlap if they have the same probability vector.

For a quantitative assessment of DeLUCS, we calculated its “classification accuracy” as defined in the Evaluation section. For the vertebrate mtDNA dataset, Table 4 shows that, at each taxonomic level, down to the genus level, our unsupervised deep learning algorithm outperforms the other two unsupervised clustering methods. Surprisingly, DeLUCS also outperforms a supervised learning algorithm that uses the same architecture in some tests (e.g., in Test 2, Class to Subclass), sometimes by a significant margin (e.g., in Test 6, Family to Genus, by 11%). Note that, in order to obtain a reasonable classification accuracy, the number *m* of mimics had to be increased for the tests/datasets with less than 150 sequences per cluster.

**Table 4.**
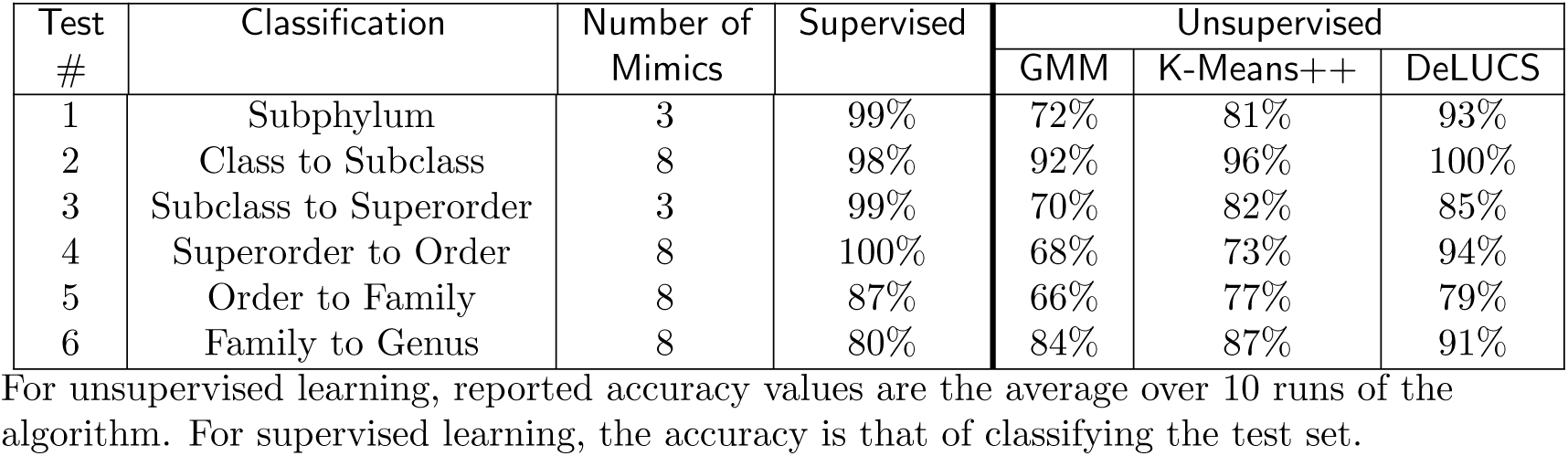
Classification accuracies for the mtDNA dataset in Table 1, Tests 1 to 6.

To test the ability of DeLUCS to cluster bacterial genome segments, and achieve a direct accuracy comparison with other unsupervised learning methods, we first attempted clustering a dataset comprising genome segments, averaging 400,000 bp, from the eight different bacterial families considered in [11] (Table 5, Test 7). The DeLUCS classification accuracy of 77% is 19% to 21% higher than the accuracies of the other two unsupervised learning methods (GMM 58%, and K-means++ 56%). The relatively low classification accuracies for all three unsupervised methods may be due in part to this dataset having a very heterogeneous evolutionary composition, with recent changes in the taxonomy of all eight bacterial families resulting during the reclassification and transition from NCBI to GTDB. In particular, *Bacillaceae/ Staphylococcaceae* and *Clostridiaceae* (formerly all Firmicutes) are now split into two different phyla, and similarly *Enterobacteriaceae/ Burkholderiaceae/ Rhodobacteraceae* and *Desulfovibrionaceae* (formerly all Proteobacteria) are now split into two different phyla. The heterogeneity of the dataset makes clustering a challenging task, as the algorithm attempts to determine cluster labels simultaneously for both closely related families and distantly related families. The patterns observed in the confusion matrix (see *S3 Appendix: Confusion matrices)* support the hypothesis of heterogeneity in genetic distance between members of the dataset. Indeed, misclassification between phyla are a minority, and most of the misclassifications occur among families that were previously placed within the same phylum, but are now placed in different phyla.

To verify that the lower classification accuracy could be partially caused by the heterogeneity of the dataset, we next considered an intra-phylum clustering of a subset of this dataset, comprising only sequences belonging to phylum Proteobacteria (Table 5, Test 8). As predicted, the classification accuracy of DeLUCS shows a significant increase, from 77% to 90%, now outperforming the other two unsupervised learning methods by 40% and 48% respectively. The majority of the misclassified genome segments in Test 8 belong to the family *Desulfobivrionaceae.* This may be partly due to this family having the shortest genome segments: The average *Desulfobivrionaceae* genome segment length is 359,337 bp, significantly shorter than the average genome segment length in Test 7, which is 433,613 bp.

**Table 5.** Classification accuracy for the bacterial datasets in Table 2, Test 7 and 8.

| Test # | Classification | Number of Mimics | Supervised | Unsupervised |  |  |
| --- | --- | --- | --- | --- | --- | --- |
|  |  |  |  | GMM | K-Means++ | DeLUCS |
| 7 | Bacteria into families | 3 | 98% | 58% | 60% | 77% |
| 8 | Proteobacteria into families | 3 | 99% | 50% | 43% | 90% |

For unsupervised learning, reported accuracy values are the average over ten runs. For supervised learning, the reported accuracy is that of classifying the test set.

Note that, in both Tests 7 and 8, we used the default value m = 3 for the number of mimic sequences. This is because the number of available sequences (genome segments of the required size) per cluster was large, and the classification accuracy did not increase by increasing the number of mimic sequences.

To test the ability of DeLUCS to cluster closely related sequences, we next attempted clustering of three viral sequence datasets, *Influenza A* virus NA-encoding gene, *Dengue* virus complete genomes, and *Hepatitis B* virus complete genomes into clusters corresponding to virus subtypes. The classification accuracies presented in Table 6 show that all machine learning methods, supervised or unsupervised, perform well, in spite of the fact that the viral sequences are very similar. In Tests 9, 10, 11 the default value m = 3 for the number of mimic sequences was used, since this was sufficient to obtain near perfect classification accuracy.

**Table 6.** Classification accuracy for the viral sequences datasets, Table 3, Tests 9, 10, and 11.

| Test # | Classification | Number of Mimics | Supervised | Unsupervised |  |  |
| --- | --- | --- | --- | --- | --- | --- |
|  |  |  |  | GMM | K-Means++ | DeLUCS |
| 9 | Influenza A | 3 | 100% | 96% | 99% | 99% |
| 10 | Dengue | 3 | 100% | 100% | 100% | 100% |
| 11 | HBV | 3 | 100% | 100% | 100% | 100% |

For unsupervised learning, reported accuracy values are the average over ten runs. For supervised learning, the reported accuracy is that of classifying the test set.

Together, these computational tests show that, in spite of the absence of any information external to the primary sequences, DeLUCS is capable of learning meaningful clusters from unlabelled, raw, and sometimes non-homologous DNA sequences. Moreover, DeLUCS outperforms classical unsupervised clustering algorithms that use k-mer counts as input features, often by a significant margin.

### Methodology Results

#### Number of mimic sequences

The default number of mimic sequences was chosen to be *m* = 3, to correspond to the three different mechanisms used to generate mimic sequences from the original sequences: transitions, transversions, or combinations of both.

For the datasets in this study it was empirically determined that, if the cluster sizes are similar, and each cluster has more than 150 sequences, then the default value of *m* = 3 results in optimal performances (as is the case in Tests 1, 3, 7, 8, 9, 10 and 11).

For the datasets with less than 150 sequences per cluster, we tested the hypothesis that the classification accuracy could be improved by generating more artificial mimic sequences per original DNA sequence. Our observations confirmed this hypothesis in Tests 2, 4, 5, and 6, where increasing the number of mimics to *m* = 8 resulted in improved classification accuracies.

Note that, for the datasets with more than 150 DNA sequences per cluster (e.g., Tests 1, 3, 7, 8), increasing the number of mimics did not always result in increased classification accuracy. Fig 6 illustrates the effect of increasing the number of mimic sequences on classification accuracy for four different tests (Tests 2, 4 and 6, with fewer than 150 sequences/cluster, and Test 3 with more than 150 sequences/cluster). Note also that Tests 9, 10, 11 had more than 150 sequences per cluster, as well as near perfect classification accuracies, and thus no increase in the number of mimic sequences was explored.

**Fig 6.**
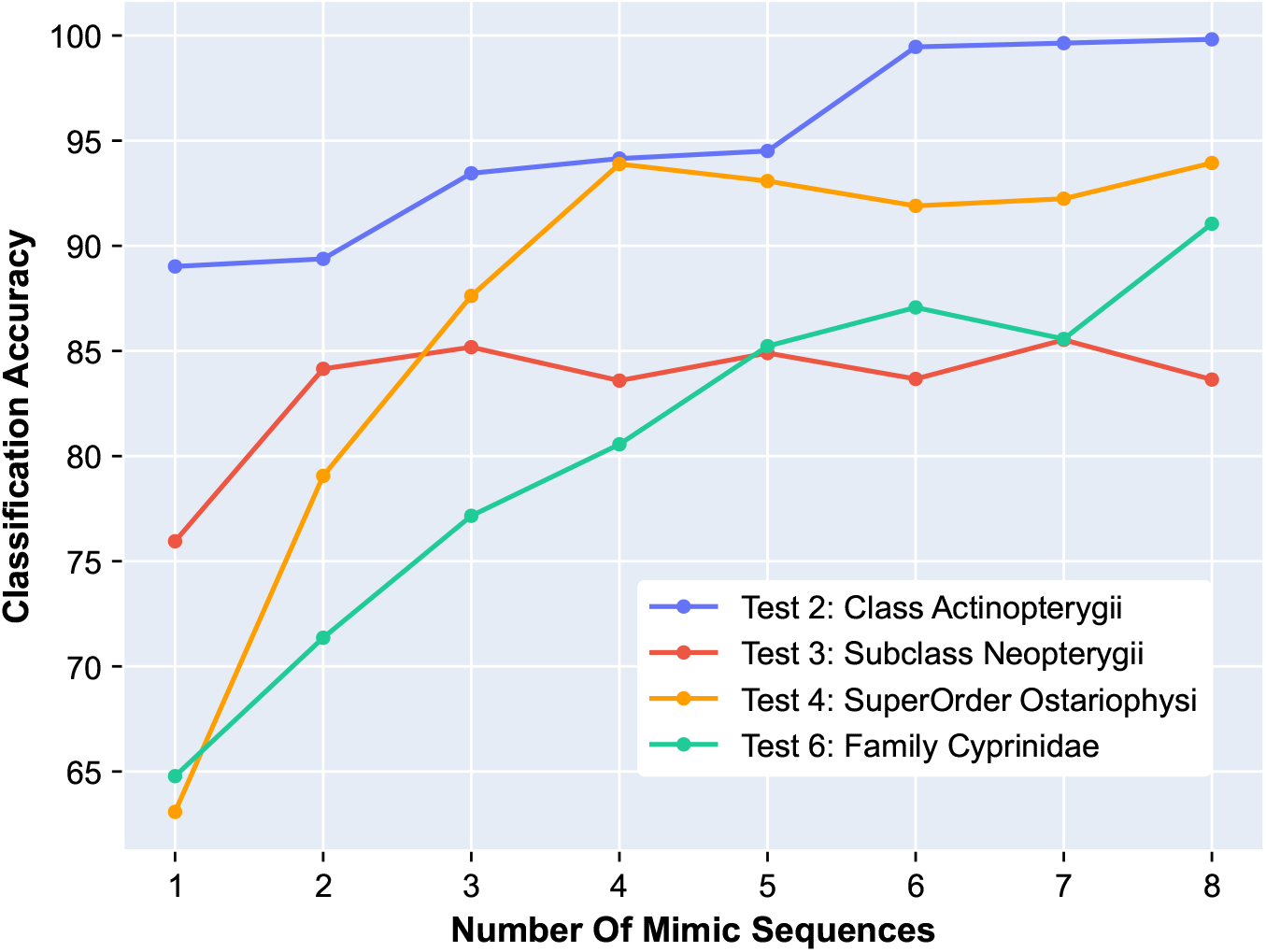
The effect of the number of mimic sequences on classification accuracy. Blue - Class Actinopterygii (Test 2); Red - Subclass Neopterygii (Test 3); Yellow - SuperOrder Ostariophysi (Test 4); Green - Family Cyprinidae (Test 6). In Tests 2, 4, and 6, with fewer than 150 sequences per cluster, increasing the number of mimics per sequence results in a marked increase in classification accuracy. In Test 3 (red), where the number of available sequences per cluster is sufficiently high (226 to 250), increasing the number of mimics to more than 3 does not result in an increase the classification accuracy.

In general, the optimal number of mimics per sequence may depend on the number of available sequences per cluster, as well as on other particulars of the dataset being analyzed. The values of m mentioned above (*m* = 3, respectively *m* = 8) are suggested, but further optimization may be possible through a hyperparameter search.

#### Adding Gaussian Noise

We now provide further insight into the performance of our method by showing the relationship between the loss function and the DeLUCS classification accuracy. The learning curves presented in Fig 7 illustrate the optimization process of a single ANN. During each epoch, the entire set of training sequences is utilized for optimizing the network parameters, and we observe an inverse correlation between the classification accuracy and the unsupervised loss function for each epoch (Fig 7, top). The same graph illustrates that the ANN sometimes tends to converge to a suboptimal solution, and this is because the mutual information might still be high for suboptimal solutions where relatively many related sequences are similarly misplaced (i.e., assigned to the wrong cluster, while still being close to each other within their subgroup).

**Fig 7.**
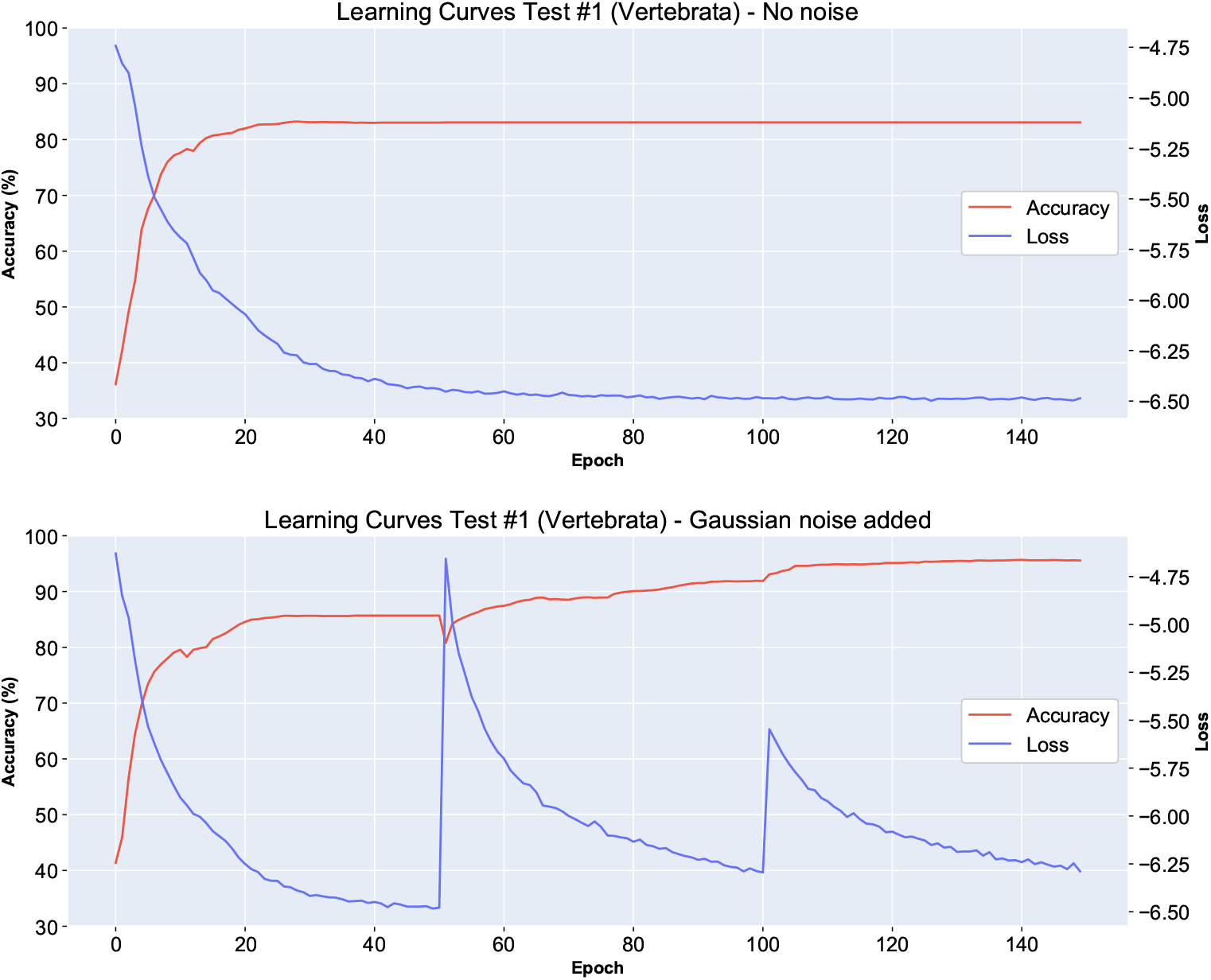
Learning curves for the training process of a single ANN in Step 2 of the Method, for the dataset in Test 1 (vertebrate mtDNA genomes) with (top), and without (bottom) the addition of Gaussian noise to the parameters of the networks every 50 epochs. Each graph displays the relation between the unsupervised loss function that is being minimized and the overall accuracy of the method. The true labels of the sequences are only used to evaluate the accuracy at each epoch, but they do not influence the learning process. The graph illustrates the positive effect that the periodic addition of Gaussian noise has on the prevention of convergence to suboptimal solutions, and on the classification accuracy (from ≈ 82% top, to ≈ 96%, bottom).

To prevent the model from converging to suboptimal solutions, we added Gaussian noise to the network parameters every 50 epochs. We confirmed empirically that the introduction of noise is beneficial, as the accuracy increases after the introduction of noise every 50 epochs (Fig 7, bottom).

#### Majority Voting for Variance Mitigation

The training process of an ANN is a randomized algorithm, with the randomness introduced by the initialization of the ANN, and the selection of random batches of training data in each epoch. The random selection of batches is beneficial for the marginalization that is performed to calculate the loss function. However, we also observed a high variance in the outcomes of training several independent copies of the ANN over the same dataset, likely due to the aforementioned randomness. To overcome this challenge, for each dataset, we independently trained ten ANNs, and integrated the obtained clustering results using a majority voting scheme. Fig 8 compares the accuracy of six tests that use an ensemble of 10 ANNs, with majority voting, compared to six tests that use a single ANN. Each test was repeated ten times to compare the variance of the two approaches. As Fig 8 shows, majority voting not only reduced the variance of predictions but also improved the overall classification accuracies.

**Fig 8.**
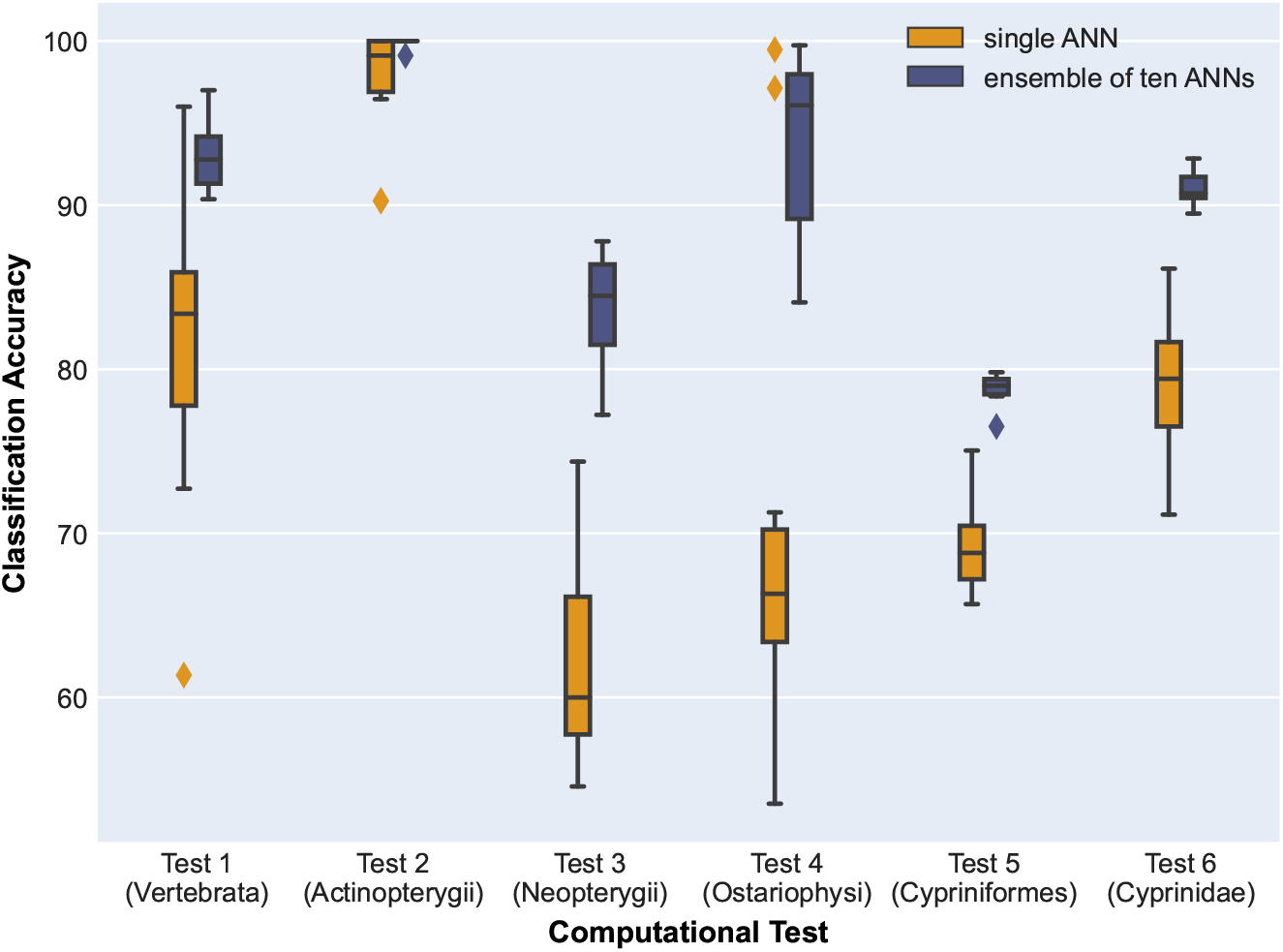
Comparison between training a single ANN (yellow), versus training an ensemble of 10 ANNs (blue) and using majority voting to decide the final cluster label for each sequence (Tests 1-6). All tests were repeated 10 times. The minimum, first quartile (Q1), median, third quartile (Q3), and maxima of the distributions are shown. The diamonds outside the boxes represent outliers. Note that the result variance for single ANNs is larger than the result variance for the corresponding ensembles of 10 ANNs. In addition, the classification accuracy is higher for the majority voting, in all cases.

## Discussion

The use of FCGR as a numerical representation of genomic sequences, in combination with invariant information clustering (a deep unsupervised learning method for computer vision), allows DeLUCS to accurately cluster large datasets of genomic sequences. The largest computational experiment in this paper comprises 3,200 randomly selected bacterial genome segments, totalling more than 1 billion bp, a dataset which is a full order of magnitude larger than previous studies clustering genomic data [8–14].

In addition, DeLUCS achieves significantly higher classification accuracies compared to other unsupervised machine learning clustering methods (K-means++ and GMM), in comparable time. The running time of the whole pipeline of DeLUCS, considering both training and testing time, is less than 40 minutes for the largest bacterial dataset

Note that for datasets comprising sequences with minimal homology (Tests 1-6, and 10-11) using alignment-based methods would be near impossible, while for datasets comprising non-homologous sequences (Tests 7, 8) using alignment-based methods would be impossible. Indeed, in the former case (vertebrate mitochondrial genomes, and virus genomes respectively), the time complexity of multiple-sequence alignment is prohibitive. In the latter case (bacterial genome segments averaging 400 kbp), the random sampling of genome segments from bacterial genomes almost guarantees non-homologous regions, thus making alignment impossible even if time complexity were not an issue. Test 9, of the NA-encoding gene of *Influenza A* virus, where an alignment-based method would be possible and probably successful, was included to showcase the applicability of our method to a variety of genomic data.

This study serves as a proof of concept of the ability of unsupervised deep learning to effectively cluster unlabelled raw DNA sequences. DeLUCS is a fully-automated method that could be used for successful clustering of datasets where traditional methods are not applicable, such as: large datasets of dissimilar and variable length raw DNA sequences, DNA sequences with incomplete or missing biological annotations, and DNA sequences for which taxonomic information or other sequence identifiers are insufficient or unavailable. Future work can address some of the current limitations of this approach, as described below.

First, in order to achieve accurate clustering, DeLUCS requires its training dataset to have balanced and well-represented clusters, with at least a minimum number of sequences per cluster. The minimum and maximum cluster size were determined individually for each computational test, based on the number of available sequences and the aforementioned requirements. As a result, many available sequences were not included in the training data, either due to the fact that under-represented clusters were discarded, or due to the fact that the excess sequences from over-represented clusters were not used. This limitation could be addressed, e.g., by adjusting the loss function to be able to deal with unbalanced training databases, that is, datasets where clusters are of different sizes.

Second, we observe that the classification accuracy of an ANN is heavily dependent on the initialization of the parameters, which is random for each run of the experiment. In other words, the classification accuracy for a dataset can vary from run to run of an ANN, sometimes by a large amount. On the other hand, one of the reasons behind DeLUCS’ successful clustering capability lies in this randomness in the parameter initialization. We attempted to address this trade-off between performance and reproducibility by training several copies of the same ANN, and using majority voting to determine the final cluster labels. The overall classification accuracy stabilized and increased as a result, with the downside being that this approach increased the running time of the training step, since ten ANNs were sequentially trained. More time-efficient solutions, such as training the ANNs in parallel, would lead to a tenfold improvement of the running time.

Third, we observed that for the datasets with fewer than 150 sequences per cluster, an increase in the number of mimics resulted in classification accuracy improvements (Tests 2, 4, 5, 6). However, this was not the case for some of the datasets with more than 150 sequences per cluster (Tests 1, 3, 7, 8). Some other tests, e.g., Tests 9, 10, 11, resulted in near perfect accuracy from the start and needed no further optimization. For experiments in this paper, the number of mimic sequences per cluster was empirically determined to be optimal with respect to the cluster size. Further exploration is needed to determine the relationship between cluster size and the number of mimics per sequence, as well as to find other mechanisms to boost classification accuracy for specific datasets, such as the use of Convolutional Neural Networks which make full use of the two-dimensional aspect of the FCGR representation.

## Conclusions

In this work we introduce DeLUCS, a novel unsupervised deep learning clustering method for DNA sequences. DeLUCS leverages deep learning to discover identity-relevant patterns in raw, primary DNA sequence data, without requiring homology, biological annotations, or the time-consuming and laborious step of defining taxonomic labels for the training data. DeLUCS obtains high classification accuracies by the novel fusion of bioinformatics approaches with recent developments in the field of deep learning for computer vision, through the use of Chaos Game Representation of original and mimic DNA sequences as input for invariant information clustering.

This is, to the best of our knowledge, the first effective alignment-free method that utilizes deep ANNs for unsupervised clustering of unlabelled DNA sequences. DeLUCS clusters diverse datasets comprising thousands of homology-free, identifier-free sequences (vertebrate mtDNA full genomes at various taxonomic levels; random segments of bacterial genomes into families; viral genomes into virus subtypes). Following the determination of the optimal mapping between the clusters learned by DeLUCS and true taxonomic groups (via the Hungarian algorithm), we note that the resulting classification is of the highest sensitivity at the species into subtypes level (99% to 100%), while in all other tests classification accuracy is double-digit higher than that of classic unsupervised machine learning clustering methods.

## Supporting information

*[S1 Appendix: Instructions for reproduction of the tests using DeLUCS](https://github.com/millanp95/DeLUCS/blob/master/paper/Supplementary/Appendix%20S1.pdf)*

*[S2 Appendix: Query options for data download](https://github.com/millanp95/DeLUCS/blob/master/paper/Supplementary/Appendix%20S2.pdf)*

*[S3 Appendix: Confusion matrices](https://github.com/millanp95/DeLUCS/blob/master/paper/Supplementary/Appendix%20S3.pdf)*

*[S4 Appendix: Detailed description of the mtDNA datasets](https://github.com/millanp95/DeLUCS/blob/master/paper/Supplementary/Appendix%20S4.pdf)*

*[S5 Appendix: A note on comparing DeLUCS with other deep learning-based clustering methodologies](https://github.com/millanp95/DeLUCS/blob/master/paper/Supplementary/Appendix%20S5.pdf)*

*[S6 Appendix: Using NMI and ARI to compare DeLUCS with K-means++ and GMM](https://github.com/millanp95/DeLUCS/blob/master/paper/Supplementary/Appendix%20S6.pdf)*

## Acknowledgments

We thank Gurjit Randhawa and Maximillian Soltysiak for suggestions on a previous version of this manuscript, and David Chen for suggestions regarding Figure 1.

## Author Contributions

**Conceptualization:** P. Millan Arias, F. Alipour, K.Hill, L.Kari

**Data curation:** P. Millan Arias, F. Alipour

**Formal analysis:** P. Millán Arias, F. Alipour, L.Kari

**Funding acquisition:** K.Hill, L. Kari

**Investigation:** P. Millán Arias, F. Alipour, K.Hill, L. Kari

**Methodology:** P. Millán Arias, F. Alipour, L. Kari

**Project administration:** L. Kari

**Resources:** K. Hill, L. Kari

**Software:** P. Millán Arias, F. Alipour

**Supervision:** L. Kari

**Validation:** P. Millán Arias, F. Alipour

**Visualization:** P. Millán Arias, F. Alipour

**Writing – original draft:** P. Millán Arias, F. Alipour

**Writing – review & editing:** P. Millán Arias, F. Alipour, K. Hill, L. Kari

