## Appendix 6 for "DeLUCS: Deep Learning for Unsupervised Clustering of DNA Sequences": Appendix 6.pdf

### S6 Appendix

#### Using NMI and ARI to compare DeLUCS with K-means++ and GMM

- **Normalized Mutual Information (NMI**). Computes and normalizes the mutual information between cluster assignments and the ground truth. It ranges from 0 (no mutual information) to 1 (perfect correlation).
- **Adjusted Rand Index (ARI).** Measures a similarity score (range: -1 to 1) with the ground truth. Random assignments have an ARI close to 0, while 1 stands for a perfect match.

**Table 1.** Comparison of DeLUCS, K-means++, and GMM, on all test datasets in the paper, using the average of the external validation measures NMI and ARI, over ten independent runs of the algorithms on each dataset. Higher is better. Boldface indicates highest/best.

| Test # | method | NMI | ARI |
|---|---|---|---|
| 1 (Vertebrata) | DeLUCS | **0.8980** | **0.9039** |
| | GMM | 0.5815 | 0.5155 |
| | *K*-Means++ | 0.6510 | 0.6330 |
| 2 (Actinopterygii) | DeLUCS | **1.0000** | **1.0000** |
| | GMM | 0.8818 | 0.8798 |
| | *K*-Means++ | 0.8617 | 0.8562 |
| 3 (Neopterygii) | DeLUCS | **0.7011** | **0.6522** |
| | GMM | 0.5312 | 0.4408 |
| | *K*-Means++ | 0.6859 | 0.5915 |
| 4 (Ostariophysi) | DeLUCS | **0.7345** | **0.7528** |
| | GMM | 0.4765 | 0.3969 |
| | *K*-Means++ | 0.5620 | 0.4867 |
| 5 (Cypriniformes) | DeLUCS | **0.7270** | **0.6008** |
| | GMM | 0.6909 | 0.5654 |
| | *K*-Means++ | 0.7264 | 0.6002 |
| 6 (Cyprinidae) | DeLUCS | 0.8846 | 0.8211 |
| | GMM | 0.8999 | 0.8273 |
| | *K*-Means++ | **0.9012** | **0.8366** |
| 7 (Bacteria) | DeLUCS | **0.7005** | **0.6117** |
| | GMM | 0.6667 | 0.5492 |
| | *K*-Means++ | 0.5854 | 0.4453 |
| 8 (Proteobacteria) | DeLUCS | **0.7723** | **0.7521** |
| | GMM | 0.4147 | 0.3212 |
| | *K*-Means++ | 0.2318 | 0.1579 |
| 9 (Influenza-A) | DeLUCS | 0.9597 | 0.9680 |
| | GMM | **0.9799** | **0.9844** |
| | *K*-Means++ | **0.9799** | **0.9844** |

| | | | |
|---|---|---|---|
| 10 (Dengue) | DeLUCS<br>GMM<br>*K*-Means++ | **1.0000**<br>**1.0000**<br>0.9999 | **1.0000**<br>**1.0000**<br>**1.0000** |
| 11 (HBV) | DeLUCS<br>GMM<br>*K*-Means++ | **1.0000**<br>0.8998<br>**1.0000** | **1.0000**<br>0.7729<br>**1.0000** |
