## Appendix 1 for "DeLUCS: Deep Learning for Unsupervised Clustering of DNA Sequences": Appendix 1.pdf

#### Instructions for the Reproduction of the Results in the Paper

First, go to https://github.com/millanp95/DeLUCS and clone the repository. Then run the series of commands for each test.

##### Test #1:

1. python build_dp.py --data_path='../data/Vertebrata/Test Files'
2. python get_pairs.py --data_path='../data/Vertebrata/Test Files/train.p' --k=6 --modify='mutation' --output='../data/Vertebrata/Test Files/testing_data.p'
3. python EvaluateDeLUCS.py --data_dir='../data/Vertebrata/Test Files' --out_dir='../data/Vertebrata/Test Files'

Comparison Models:

- python EvaluateComparison.py --data_path='../data/Vertebrata/Test Files/testing_data.p'  --method='Supervised' --k=6
- python EvaluateComparison.py --data_path='../data/Vertebrata/Test Files/train.p'  --method='k-means++' --k=6
- python EvaluateComparison.py --data_path='../data/Vertebrata/Test Files/train.p'  --method='GMM' --k=6 --k=6

##### Test #2:

1. python build_dp.py --data_path='../data/Fish/Test Files/Actinopterygii'
2. python get_pairs.py --data_path='../data/Fish/Test Files/Actinopterygii/train.p' --k=6 --n_mimics=8 --modify='mutation' --output=../data/Fish/Test Files/Actinopterygii/testing_data.p'
3. python EvaluateDeLUCS.py --data_dir='../data/Fish/Test Files/Actinopterygii' --out_dir='../data/Fish/Test Files/Actinopterygii'

Comparison Models:

- python EvaluateComparison.py --data_path='../data/Fish/Test Files/Actinopterygii/testing_data.p'   --method='Supervised' --k =6
- python EvaluateComparison.py --data_path='.../data/Fish/Test Files/Actinopterygii/train.p'   --method='k-means++' --k=6
- python EvaluateComparison.py --data_path='../data/Fish/Test Files/Actinopterygii/train.p'   --method='GMM' --k=6

**Test #3:**

1. python build_dp.py --data_path='../data/Fish/Test Files/Neopterygii'
2. python get_pairs.py --data_path='../data/Fish/Test Files/Neopterygii/train.p' --k=6 --modify='mutation' --output=../data/Fish/Test Files/Neopterygii/testing_data.p'
3. python EvaluateDeLUCS.py --data_dir='../data/Fish/Test Files/Neopterygii' --out_dir='../data/Fish/Test Files/Neopterygii'

   Comparison Models:

- python EvaluateComparison.py --data_path='../data/Fish/Test Files/Neopterygii/testing_data.p'  --method='Supervised' --k =6
- python EvaluateComparison.py --data_path='../data/Fish/Test Files/Neopterygii/train.p'  --method='k-means++' --k=6
- python EvaluateComparison.py --data_path='../data/Fish/Test Files/Neopterygii/train.p'  --method='GMM' --k=6

**Test #4:**

1. python build_dp.py --data_path='../data/Fish/Test Files/Ostariophysi'
2. python get_pairs.py --data_path='../data/Fish/Test Files/Ostariophysi/train.p' --k=6 --n_mimics=8 --modify='mutation' --output=../data/Fish/Test Files/Ostariophysi/testing_data.p'
3. python EvaluateDeLUCS.py --data_dir='../data/Fish/Test Files/Ostariophysi' --out_dir='../data/Fish/Test Files/Ostariophysi'

   Comparison Models:

- python EvaluateComparison.py --data_path='./data/Fish/Test Files/Ostariophysi/testing_data.p' --method='Supervised' --k =6
- python EvaluateComparison.py --data_path='../data/Fish/Test Files/Ostariophysi/train.p' --method='k-means++' --k=6
- python EvaluateComparison.py --data_path='../data/Fish/Test Files/Ostariophysi/train.p'  --method='GMM' --k=6

**Test #5:**

1. python build_dp.py --data_path='../data/Fish/Test Files/Cypriniformes'
2. python get_pairs.py --data_path='../data/Fish/Test Files/Cypriniformes/train.p' --k=6 --n_mimics=8 --modify='mutation' --output=../data/Fish/Test Files/Cypriniformes/testing_data.p'
3. python EvaluateDeLUCS.py --data_dir='../data/Fish/Test Files/Cypriniformes' --out_dir='../data/Fish/Test Files/Cypriniformes'

   Comparison Models:

- python EvaluateComparison.py --data_path='../data/Fish/Test Files/Cypriniformes/testing_data.p'  --method='Supervised' --k =6
- python EvaluateComparison.py --data_path='../data/Fish/Test Files/Cypriniformes/train.p' --method='k-means++' --k=6
- python EvaluateComparison.py --data_path='../data/Fish/Test Files/Cypriniformes/train.p' --method='GMM' --k=6

#### **Test #6:**

1. python build_dp.py --data_path='../data/Fish/Test Files/Cyprinidae'
2. python get_pairs.py --data_path='../data/Fish/Test Files/Cyprinidae/train.p' --k=6 --n_mimics=8 --modify='mutation' --output=../data/Fish/Test Files/Cyprinidae/testing_data.p'
3. python EvaluateDeLUCS.py --data_dir='../data/Fish/Test Files/Cyprinidae' --out_dir='../data/Fish/Test Files/Cyprinidae'

<u>Comparison Models:</u>

- python EvaluateComparison.py --data_path='../data/Fish/Test Files/Cyprinidae/testing_data.p'   --method='Supervised' --k =6
- python EvaluateComparison.py --data_path='../data/Fish/Test Files/Cyprinidae/train.p'  --method='k-means++' --k=6
- python EvaluateComparison.py --data_path='../data/Fish/Test Files/Cyprinidae/train.p' --method='GMM' --k=6

#### **Test #7:**

1. python build_dp.py --data_path='../data/Bacteria/Test Files'
2. python get_pairs.py --data_path='../data/Bacteria/Test Files/train.p' --k=6 --modify='mutation' --output='../data/Bacteria/Test Files/testing_data.p'
3. python EvaluateDeLUCS.py --data_dir='../data/Bacteria/Test Files' --out_dir='../data/Bacteria/Test Files'

<u>Comparison Models:</u>

- python EvaluateComparison.py --data_path='../data/Bacteria/Test Files/testing_data.p'  --method='Supervised' --k =6
- python EvaluateComparison.py --data_path='../data/Bacteria/Test Files/train.p' --method='k-means++' --k=6
- python EvaluateComparison.py --data_path='../data/Bacteria/Test Files/train.p' --method='GMM' --k=6

#### **Test #8:**

1. python build_dp.py --data_path='../data/ProteoBacteria/Test Files'
2. python get_pairs.py --data_path='../data/ProteoBacteria/Test Files/train.p' --k=6 --modify='mutation' --output='../data/ProteoBacteria/Test Files/testing_data.p'
3. python EvaluateDeLUCS.py --data_dir='../data/ProteoBacteria/Test Files' --out_dir='../data/ProteoBacteria/Test Files'

<u>Comparison Models:</u>

- python EvaluateComparison.py --data_path='../data/ProteoBacteria/Test Files/testing_data.p' --method='Supervised' --k =6
- python EvaluateComparison.py --data_path='../data/ProteoBacteria/Test Files/train.p' --method='k-means++' --k=6
- python EvaluateComparison.py --data_path='../data/ProteoBacteria/Test Files/train.p' --method='GMM' --k=6

#### Test #9:

1. python build_dp.py --data_path='../data/Influenza-A/Test Files'
2. python get_pairs.py --data_path='../data/Influenza-A/Test Files/train.p' --k=6 --modify='mutation' --output='../data/Influenza-A/Test Files/testing_data.p'
3. python EvaluateDeLUCS.py --data_dir='../data/Influenza-A/Test Files' --out_dir='../data/Influenza-A/Test Files'

   Comparison Models:

- python EvaluateComparison.py --data_path='../data/Influenza-A/Test Files/testing_data.p'  --method='Supervised' --k =6
- python EvaluateComparison.py --data_path='../data/Influenza-A/Test Files/train.p' --method='k-means++' --k=6
- python EvaluateComparison.py --data_path=../data/Influenza-A/Test Files/train.p' --method='GMM' --k=6

#### Test #10:

1. python build_dp.py --data_path='../data/Dengue/Test Files'
2. python get_pairs.py --data_path='../data/Dengue/Test Files/train.p' --k=6 --modify='mutation' --output='../data/Dengue/Test Files/testing_data.p'
3. python EvaluateDeLUCS.py --data_dir='../data/Dengue/Test Files' --out_dir='../data/Dengue/Test Files'

   Comparison Models:

- python EvaluateComparison.py --data_path='../data/Dengue/Test Files/testing_data.p' --method='Supervised' --k =6
- python EvaluateComparison.py --data_path='../data/Dengue/Test Files/train.p' --method='k-means++' --k=6
- python EvaluateComparison.py --data_path='../data/Dengue/Test Files/train.p' --method='GMM' --k=6

#### Test #11:

1. python build_dp.py --data_path='../data/HBV/Test Files'
2. python get_pairs.py --data_path='../data/HBV/Test Files/train.p' --k=6 --modify='mutation' --output='../data/HBV/Test Files/testing_data.p'
3. python EvaluateDeLUCS.py --data_dir='../data/HBV/Test Files' --out_dir='../data/HBV/Test Files'

   Comparison Models:

1. python EvaluateComparison.py --data_path='../data/HBV/Test Files/testing_data.p'  --method='Supervised' --k =6
2. python EvaluateComparison.py --data_path='../data/HBV/Test Files/train.p' --method='k-means++' --k=6
3. python EvaluateComparison.py --data_path='../data/HBV/Test Files/train.p' --method='GMM' --k=6

### DeLUCS computational pipeline for running your own dataset:

1. Build the dataset:

```
python build_dp.py --data_path=<PATH_sequence_folder>
```

   - Input: Folders with the sequences in FASTA format
   - Output : file in the form (label,sequence,accession)

If the true label is unknown then place ALL the sequences in the same folder

2. Compute the mimics.

```
 python get_pairs.py --data_path=<PATH_dataset> --k=6 --modify='mutation'
--output=<PATH_output_file>
```

   - Input: file in the form (label, sequence, accession)
   - Output : file in the form of (pairs, x_test, y_test).

3. Train the model.

If the true labels are unknown for your dataset and you want to use DeLUCS as a clustering tool without the assessment by means of the Hungarian algorithm, run:

```
python TrainDeLUCS.py --data_dir=<PATH_of_computed_mimics> --out_dir=<OUTPURDIR>
```

   - Input: file in the form of (pairs, x_test).
   - Output : Cluster assignment for each sequence in x_test.

If the "ground truth" is known for your dataset and you want to train DeLUCS and then evaluate its performance by means of the Hungarian algorithm, run:

```
python EvaluateDeLUCS.py --data_dir=<PATH_computed_mimics> --out_dir=<OUTPURDIR>
```

   - Input: file in the form of (pairs, x_test, y_test).
   - Output : Image with the confusion matrix.
