## Appendix 2 for "DeLUCS: Deep Learning for Unsupervised Clustering of DNA Sequences": Appendix 2.pdf

### Query Options for the Download of the Original Versions of the Datasets

**Mitochondrial dataset (Nov 16, 2020) :**
We used the software Geneious and the following keywords: *Mitochondrion*, *Mitochondria*, *Vertebrata* and *Complete Genome*.

**Bacterial dataset (Jan 18, 2021):**
We downloaded the file `bac120_taxonomy_r95.tsv` directly from GTDB https://data.gtdb.ecogenomic.org/releases/release95/95.0/ and filtered the families from the complete taxonomy.

**Influenza A virus dataset (Oct 14, 2020):**
The sequences were downloaded directly from:
https://www.ncbi.nlm.nih.gov/genomes/FLU/dataset/nph-select.cgi#mainform, using the following query options:

- sequence type: Nucleotide
- type: A
- subtypes: H1N1,H2N2, H5N1, H7N3, and H7N9
- segment: 6(NA)
- other options: default
- full length only
- collapse identical

-----------Dataset Statistics ------------
Total num of classes: 5
Total num of sequences: 13078
Min genome length: 52
Avg genome length: 1392.2
Max genome length: 1544
Data distribution:

| | | |
|---|---|---|
| H5N1 | => | 3095 |
| H2N2 | => | 175 |
| H1N1 | => | 9189 |
| H7N9 | => | 293 |
| H7N3 | => | 326 |

**Dengue virus dataset (Oct 14, 2020):**
The sequences were downloaded directly from:
https://www.ncbi.nlm.nih.gov/genomes/VirusVariation/dataset/nph-select.cgi?taxid=12637, using the following query options:

- sequence type: Nucleotide
- other options: default
- collapse identical
- full-length only

-----------Dataset Statistics ------------
Total num of classes: 5
Total num of samples: 5868
Min genome length: 10161
Avg genome length: 10582.002044989775
Max genome length: 11195
Data distribution:
Subtype-1 => 2446
Subtype-2 => 1891
Subtype-3 => 1121
Subtype-4 => 407
N/A => 3

**Hepatitis B virus dataset (Oct 14, 2020):**
The whole dataset was downloaded directly from:
https://hbvdb.lyon.inserm.fr/HBVdb/HBVdbDataset?seqtype=0.

-----------Dataset Statistics ------------
Total num of classes: 6
Total num of samples: 6493
Min genome length: 3182
Avg genome length: 3209.943015555213
Max genome length: 3254
Data distribution:
A => 880
B => 1765
C => 2194
D => 1090
E => 306
F => 258

**Note:** The curated datasets used to obtain the results in the paper can be found at:
https://github.com/millanp95/DeLUCS/tree/master/data
