## Appendix 3 for "DeLUCS: Deep Learning for Unsupervised Clustering of DNA Sequences": Appendix 3.pdf

### Confusion Matrices for a Single Run of Every Computational Test

**Figure 1 (Tests 1-6).** Confusion matrices of the assignment that maximizes the accuracy at each taxonomic level of the mtDNA dataset. Predicted labels are numeric cluster assignments, omitted here for readability.

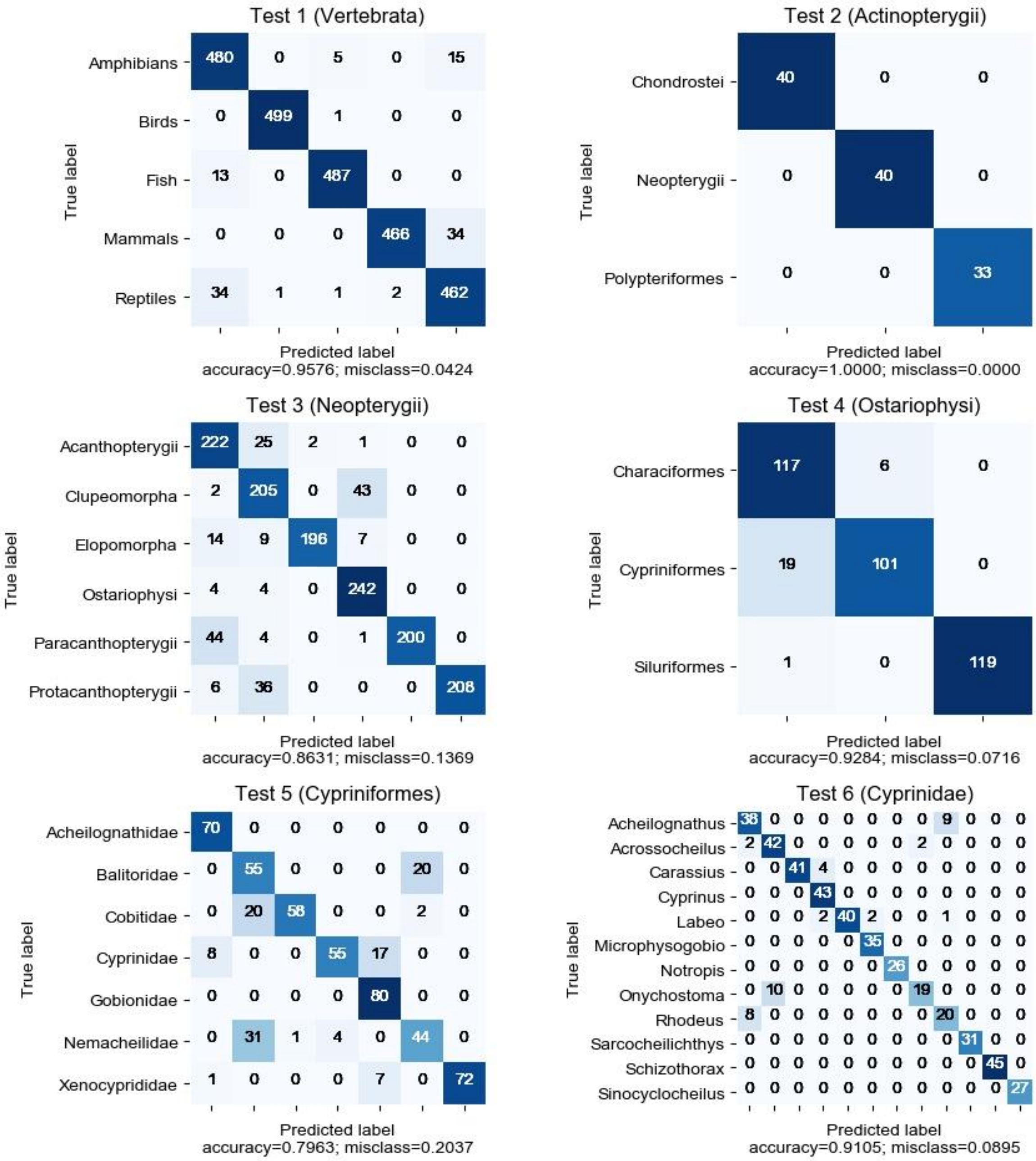

**Figure 2 (Tests 7, 8).** Confusion matrices of the assignment that maximizes the accuracy for both computational tests with the bacterial dataset at phylum level to families. (Top) All bacterial families are considered. (Bottom) Only sequences in the phylum Proteobacteria are considered. Predicted labels are numeric cluster assignments, omitted here for readability.

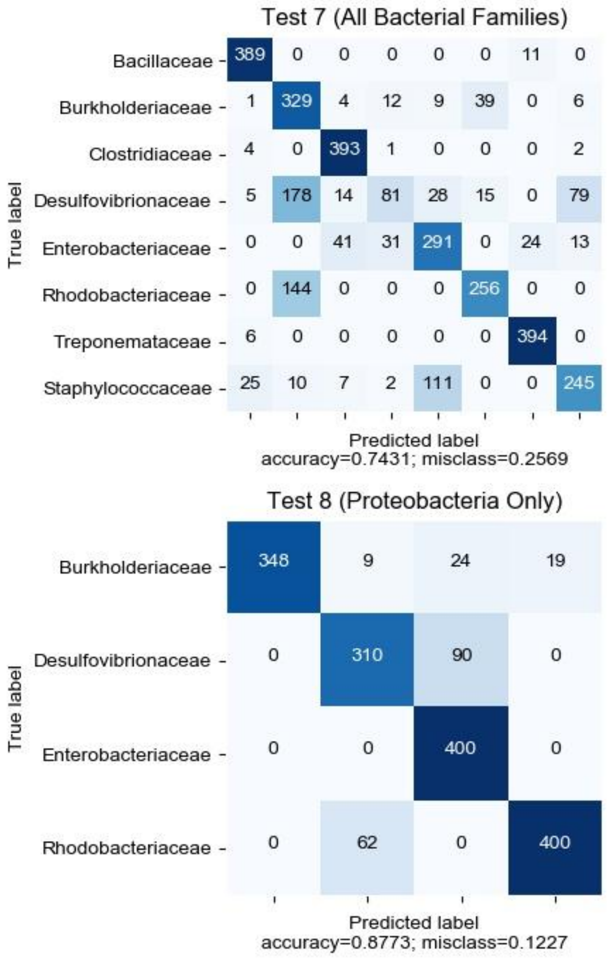

**Figure (Tests 9-11).** Confusion matrices of the assignment that maximizes the accuracy for the NA-encoding gene of the Influenza A virus, Dengue virus genomes, and HBV genomes. Predicted labels are numeric cluster assignments, omitted here for readability.

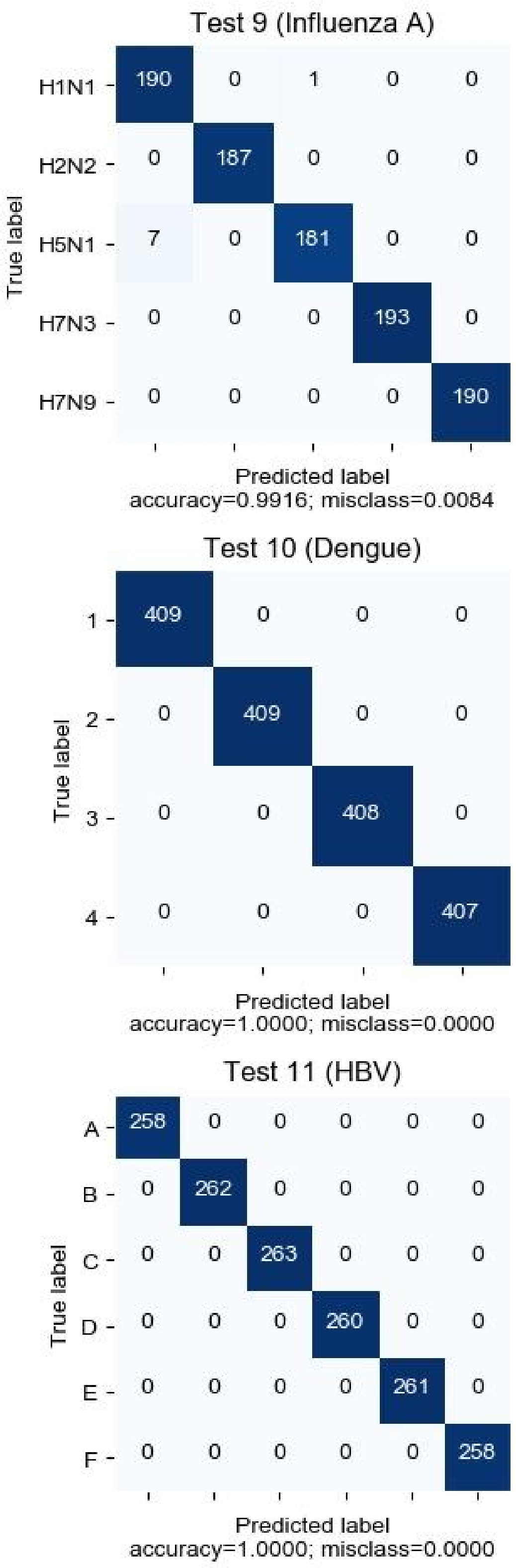
