## Appendix 4 for "DeLUCS: Deep Learning for Unsupervised Clustering of DNA Sequences": Appendix 4.pdf

### Details of the Full Vertebrate Mitochondrial Genomes Datasets (Tests 1 - 6)

**Figure 1 (mtDNA Decision Tree).** Decision tree demonstrating the cluster choices at different taxonomic levels in mtDNA dataset. The choice of computational Tests 1 to 6 follows a decision-tree approach whereby, at each level, one particular cluster is selected for further in-depth exploration. The taxonomic labels of the clusters are unknown during training and are not used by DeLUCS.

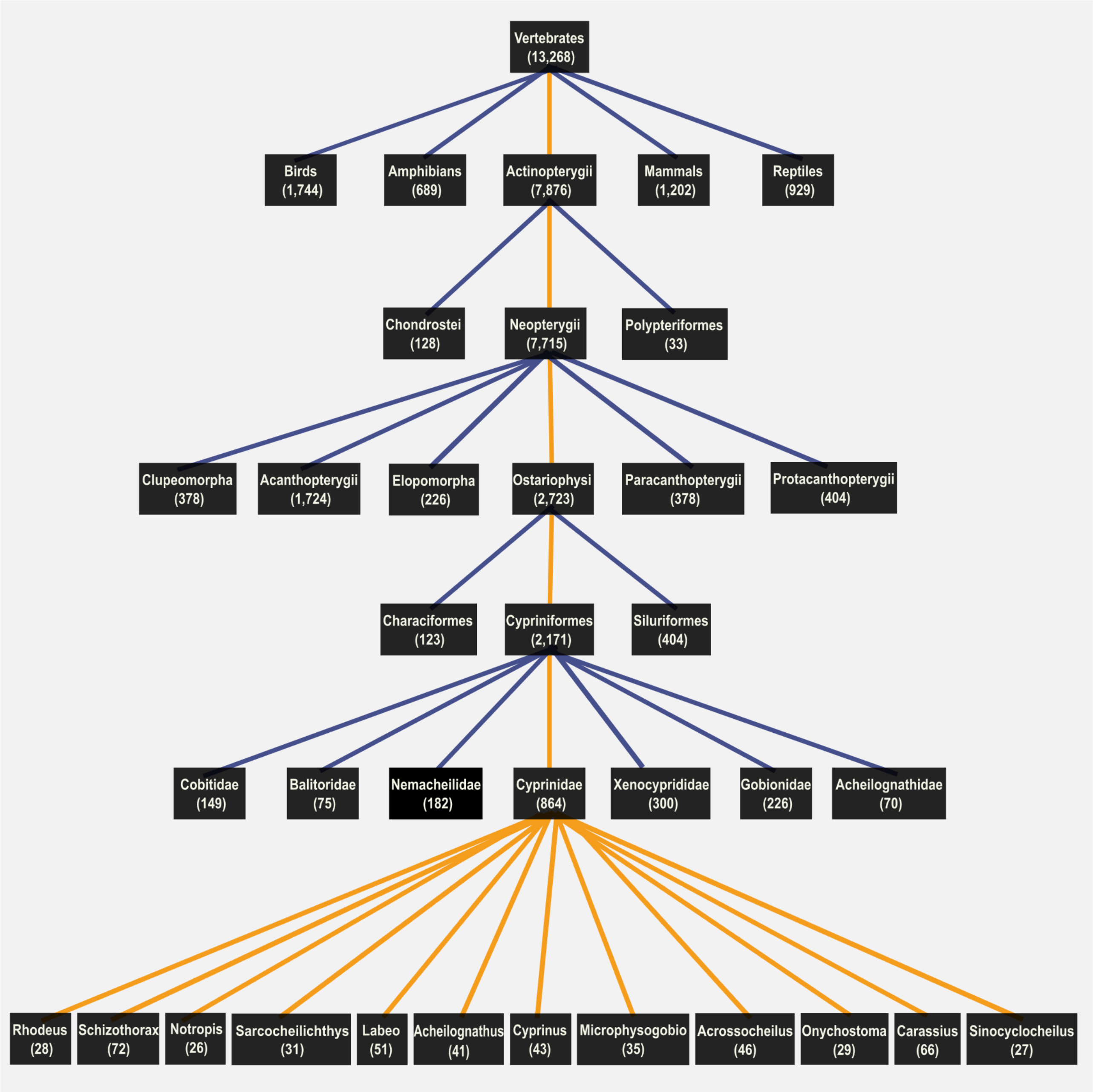

### Description of corresponding NCBI datasets

For each test: Clusters with fewer sequences than the minimum cluster size are excluded as well as the taxonomic groups without the particular identifier. This document includes the number of the sequences available on NCBI under a particular taxonomic identifier as of Nov 2020.

**Test #1: Subphylum Vertebrata**

Actinopterygii (Fish) [7876]
Amphibia [689]
Aves [1744]
Mammalia (No Humans) [1202]
Reptiles (Crocodylia [44] , Sphenodontia [36] , Squamata [650] , Testudines [199])

**Test #2: SuperClass Actinopterygii**

Chondrostei [128]
Polypteriformes [33]
Neopterygii [7715]

**Test #3: Subclass Neopterygii**

Clupeomorpha [378]
Ostariophysi [2723]
Protacanthopterygii [404]
Acanthopterygii [1724]
Paracanthopterygii [378]
Elopomorpha [226]

**Excluded clusters (super-order)**

Osteoglossomorpha [73]
Cyclosquamata [23]
Lampridiomorpha [6]
Polymyxiomorpha [4]
Scopelomorpha [12]
Stenopterygii [12]

**Excluded clades that do not have the super-order identifier**

Eupercaria [650]
Batrachoidaria [3]
Gobiaria [342]
Pelagiaria [173]
Carangaria [139]
Ovalentaria [306]
Anabantaria [67]
Zeiogadaria [3]
Otocephala [27]

**Test #4: Super-Order Ostariophysi**

Characiformes [123]
Cypriniformes [2171]

Siluriformes [404]

**Excluded clusters (order)**
Gonorynchiformes [10]
Gymnotiformes [22]

**Test #5: Order Cypriniformes**
Cyprinidae [864]
Balitoridae [75]
Nemacheilidae [182]
Cobitidae [149]
Xenocyprididae [300]
Gobionidae [226]
Acheilognathidae [70]

**Excluded clusters (family)**
Gastromyzontidae [54]
Danionidae [58]
Botiidae [42]
Psilorhynchidae [4]
Catostomidae [45]
Cyprinoidei intergeneric hybrids [58]
Tincidae [2]
Gyrinocheilidae [4]
Vaillantellidae [2]
Serpenticobitidae [1]
Leptobarbidae [2]
Tanichthyidae [2]
Barbuccidae [1]
Paedocyprididae [6]
Ellopostomatidae [1]

**Excluded sub-families that do not have the family identifier**
Leuciscidae [146]

**Test #6: Family Cyprinidae**
Rhodeus [28]
Schizothorax [72]
Notropis [26]
Carassius [66]
Labeo [51]
Microphysogobio [35]
Acrossocheilus [46]
Onychostoma [29]
Sarcocheilichthys [31]
Acheilognathus [47]
Sinocyclocheilus [27]
Cyprinus [43]

**Excluded clusters (genus)**
Danio [13]
Opsariichthys [16]
Barbus [4]
Puntius [10]
Pethia [3]
Sahyadria [4]
Ptychobarbus [9]
Diptychus [4]
Schizopygopsis [23]
Gymnocypris [20]
Hampala [2]
Herzensteinia [2]
Chuanchia [2]
Tor [19]
Gymnodiptychus [10]
Cyprinion [1]
Platypharodon [3]
Oxygymnocypris [2]
Scaphiodonichthys [2]
Mystacoleucus [2]
Luciobarbus [5]
Osteobrama [3]
Onychostoma [1]
Neolissochilus [5]
Procypris [5]
Aspiorhynchus [3]
Spinibarbus [9]
Systomus [1]
Hypselobarbus [2]
Percocypris [3]
Schizopyge [7]
Carassioides [2]
Cyclocheilichthys [2]
Labeobarbus [1]
Barbodes [2]
Caecobarbus [2]
Poropuntius [1]
Puntigrus [2]

**Excluded tribes that do not have the genus identifier**
Garrini [19]
Labeonini [16]

**Excluded clades that do not have the genus identifier**
Osteochilini [9]
Semilabeonini [34]
