## Appendix 5 for "DeLUCS: Deep Learning for Unsupervised Clustering of DNA Sequences": Appendix 5.pdf

### S5 Appendix

#### A note on comparing DeLUCS with other deep learning-based clustering methodologies

This Appendix provides further support of lines 325-332 in the manuscript justifying our selection of K-means++ and GMM as the only algorithms to be compared with DeLUCS, at the exclusion of other deep learning-based clustering methodologies. Several deep learning-based clustering approaches have been proposed in the literature, mostly for computer vision tasks, and some of these methodologies have been successfully applied to various fields in bioinformatics, see [15] for a review. However, most of these methods have not been optimized to work with DNA sequences, and we hypothesized that they would perform poorly on DNA sequence datasets. To test this hypothesis, we explored the methods listed in [15] for potential comparisons with DeLUCS.

Since most of the methods in [15] share the same working principle, and their performance over the different computer vision datasets (MNIST, REUTERS, STL-10) is comparable, we selected two representative methods and compared their performance with DeLUCS. Table 1 compares DeLUCS's clustering accuracy with that of Deep Embedded Clustering (DEC), and improved Deep Embedded Clustering (iDEC), the latter run with their default architecture and parameters and with CGRs of DNA sequences as input (average over 10 runs). As expected, and as seen in Table 1, DeLUCS outperforms both DEC and iDEC by more than 65% in some tests.

| Test # | DEC | iDEC | DeLUCS |
|---|---|---|---|
| 1 | 48% | 49% | 94% |
| 2 | 30% | 31% | 100% |
| 3 | 57% | 58% | 85% |
| 4 | 28% | 30% | 94% |
| 5 | 66% | 66% | 79% |
| 6 | 74% | 79% | 91% |
| 7 | 43% | 44% | 77% |
| 8 | 38% | 39% | 90% |
| 9 | 48% | 49% | 99% |
| 10 | 39% | 39% | 100% |
| 11 | 57% | 58% | 100% |

**Table 1:** Comparison of unsupervised clustering accuracy of two deep learning based clustering methods (DEC and iDEC) with DeLUCS

We suspect that similar results would be obtained for the other methodologies in [15], as they share the same working principle. This being said, we did not include the results in Table 1 in the main body of the manuscript as we believe that, without hyperparameter optimization and architecture engineering, this would not be a meaningful comparison. Indeed, a direct (fair) comparison of DeLUCS with the methods included in [15] would require optimizing those methods' architectures and hyperparameters to work with DNA sequences, which is a non-trivial task as well as outside the scope of this paper.

For these reasons, we opted for a comparison of the performance of DeLUCS with the K-means++ and GMM algorithms only, as they have been previously used with DNA sequence datasets and they performed reasonably well on the datasets in this paper.

**<u>To reproduce the results:</u>**

1. Clone DeLUCS repository to download the data.

```
git clone https://github.com/millanp95/DeLUCS DeLUCS
cd DeLUCS
```

2. Build the data sets.

```
python build_dp.py --data_path='../data/Vertebrata/Test Files'
python build_dp.py --data_path='../data/Fish/Test Files/Actinopterygii'
python build_dp.py --data_path='../data/Fish/Test Files/Neopterygii'
python build_dp.py --data_path='../data/Fish/Test Files/Ostariophysi'
python build_dp.py --data_path='../data/Fish/Test Files/Cypriniformes'
python build_dp.py --data_path='../data/Fish/Test Files/Cyprinidae'
python build_dp.py --data_path='../data/Bacteria/Test_Files'
python build_dp.py --data_path='../data/Bacteria/Proteo_Test_Files'
python build_dp.py --data_path='../data/Influenza-A/Test Files'
python build_dp.py --data_path='../data/Dengue/Test Files'
python build_dp.py --data_path='../data/HBV/Test Files'
```

3. Clone the original DEC/iDEC repository inside the DeLUCS folder

```
git clone https://github.com/XifengGuo/IDEC.git iDEC
```

4. Replace the old files "datasets.py", “DEC.py” and “IDEC.py” by the new ones incorporating the code to indicate that the inputs are DNA sequences and the code for the computation of the CGR for each sequence.

```
sudo chmod +x DeepComparison/replaceFiles.sh
./DeepComparison/replaceFiles.sh
```

5. Run the MNIST code to verify everything was correctly installed

```
cd iDEC
python IDEC.py mnist
```

You should get as output something along these lines:

```
acc: 0.8795285714285714
clustering time: 378 seconds.
```

6. Once you tested the installation, run the experiments with the following commands:

Test #1

```
python DEC.py cgr --data_path='../data/Vertebrata/Test Files/train.p' --
n_clusters=5
python IDEC.py cgr --data_path='../data/Vertebrata/Test Files/train.p' --
n_clusters=5
```

Test #2

```
python DEC.py cgr --data_path='../data/Fish/Test Files/Actinopterygii/train.p' --
n_clusters=3
python IDEC.py cgr --data_path='../data/Fish/Test Files/Actinopterygiitrain.p' --
n_clusters=3
```

Test #3

```
python DEC.py cgr --data_path='../data/Fish/Test Files/Neopterygii/train.p' --
n_clusters=6
python IDEC.py cgr --data_path='../data/Fish/Test Files/Neopterygii/train.p' --
n_clusters=6
```

Test #4

```
python DEC.py cgr --data_path='../data/Fish/Test Files/Ostariophysi/train.p' --
n_clusters=3
python IDEC.py cgr --data_path='../data/Fish/Test Files/Ostariophysi/train.p' --
n_clusters=3
```

Test #5

```
python DEC.py cgr --data_path='../data/Fish/Test Files/Cypriniformes/train.p' --
n_clusters=7
python IDEC.py cgr --data_path='../data/Fish/Test Files/Cypriniformes/train.p' --
n_clusters=7
```

Test #6

```
python DEC.py cgr --data_path='../data/Fish/Test Files/Cyprinidae/train.p' --
n_clusters=12
python IDEC.py cgr --data_path='../data/Fish/Test Files/Cyprinidae/train.p' --
n_clusters=12
```

Test #7

```
python DEC.py cgr --data_path='../data/Bacteria/Test_Files/train.p' --n_clusters=8
python IDEC.py cgr --data_path='../data/Bacteria/Test_Files/train.p' --n_clusters=8
```

Test #8

```
python DEC.py cgr --data_path='../data/Bacteria/Proteo_Test_Files/train.p' --
n_clusters=4
python IDEC.py cgr --data_path='../data/Bacteria/Proteo_Test_Files/train.p' --
n_clusters=4
```

Test #9

```
python DEC.py cgr --data_path='../data/Influenza-A/Test Files/train.p' --
n_clusters=4
python IDEC.py cgr --data_path='../data/Influenza-A/Test Files/train.p' --
n_clusters=4
```

Test #10

```
python DEC.py cgr --data_path='../data/Dengue/Test Files/train.p' --n_clusters=5
python IDEC.py cgr --data_path='../data/Dengue/Test Filestrain.p' --n_clusters=5
```

Test #11

```
python DEC.py cgr --data_path='../data/HBV/Test Files/train.p' --n_clusters=6
python IDEC.py cgr --data_path='../data/HBV/Test Files/train.p' --n_clusters=6
```
